# Preventing surgery induced immune suppression and metastases by inhibiting PI3K-gamma signalling in Myeloid-Derived Suppressor Cells

**DOI:** 10.1101/2024.09.08.611916

**Authors:** Leonard Angka, Gayashan Tennakoon, David P. Cook, Andre B. Martel, Marisa Market, Christiano Tanese de Souza, Emma Cummins, Ismael Samudio, Natasha Kekre, Michele Ardolino, Barbara Vanderhyden, Michael A. Kennedy, Rebecca C. Auer

**Affiliations:** Cancer Therapeutics Program, Ottawa Hospital Research Institute; Ottawa, Ontario, Canada; Department of Biochemistry, Microbiology and Immunology, University of Ottawa; Ottawa, Ontario, Canada; Department of Cellular and Molecular Medicine, University of Ottawa; Ottawa, Ontario, Canada; Division of General Surgery, Department of Surgery, University of Ottawa; Ottawa, Ontario, Canada; adMare Bioinnovations; Vancouver, British Columbia, Canada; Center for Infection, Immunity and Inflammation, University of Ottawa; Ottawa, Ontario, Canada

## Abstract

Myeloid derived suppressor cells (MDSCs) have a dominating presence in the postoperative period and mediate the suppression of Natural Killer (NK) cells and promotion of cancer metastases after surgery. However, their functional characteristics and effect on cellular immunity after surgery have not been comprehensively investigated. Here, we characterize the expansion of surgery-induced (sx) MDSCs via multi-colour flow cytometry, single-cell RNA sequencing, and functional *ex vivo* NK cell suppression assays. We then screened a small molecule library using our sx-MDSC:NK cell suppression assay to identify compounds that could inhibit sx-MDSCs. These studies provide evidence that PI3K-γ signalling is upregulated in sx-MDSCs and blockade with PI3K-γ specific inhibitors attenuates NK cell suppression in humans and mice and reduces postoperative metastases in murine models. Upregulated PI3K-γ in sx-MDSCs is a potential pathway amenable to therapeutic targeting in the postoperative period.

**One Sentence Summary:** The suppressive mechanisms of surgery-induced myeloid derived suppressor cells use PI3K signalling and are amenable to PI3K-gamma specific inhibitors.

## INTRODUCTION

Postoperative immune suppression after surgery can negatively impact a patient’s susceptibility to infections, quality of life during recovery, and likelihood of metastatic relapse (*1*). During surgery, the disruption of endothelial cell barriers causes the release of damage-associated molecular patterns (DAMPs), or “alarmins”, to alert and mobilize immune cells to the site of surgery to remove damaged cells and protect from foreign pathogens (*2, 3*). Monocytes that are recruited immediately from bone marrow reserves, release inflammatory cytokines such as IL-1, IL-6 and TNF-ɑ. While monocytes and neutrophils are the main contributors to the initial proinflammatory response, they are also critical for its resolution (*3*). To compensate for the large egress of monocytes, the body activates emergency myelopoiesis which results in a rapid release of immature myeloid cells (*4*).

These immature myeloid cells are anti-inflammatory and immunosuppressive-like cells with altered cytokine profiles (IL-10, TGF-β secretion) and reduced antigen presentation (downregulation of HLA-DR). In fact, they phenotypically and functionally resemble myeloid derived suppressor cells (MDSCs) and have been implicated in mediating postoperative immunosuppression (*5–9*). The impact of these immature myeloid cells on recovery from surgery and sepsis have been studied, but very little is known about their effect on recurrence following cancer surgery.

The central role that the immune system, particularly Natural Killer (NK) cells (*10*), plays in eradication of metastatic disease is well recognized and MDSCs have been implicated as a major player. The magnitude of immune suppression following surgery is several folds higher than immune suppression induced by the cancer itself or other treatments (*11*), such as chemotherapy, and this is a predictor of metastatic cancer recurrences (*9, 12, 13*). In murine studies, our group has established that NK cell and T cell suppression in the early postoperative period is responsible for cancer recurrence and metastases (*14–18*). We have previously confirmed that these “surgery-induced” (sx)-MDSCs are responsible for postoperative NK and T cell suppression (*17*).

In order to alleviate postoperative immune suppression by sx-MDSCs, we must first characterize the cells involved. Generally, MDSCs are described as being either polymorphonuclear (PMN-), monocytic (M-), or early stage (E-) MDSCs (*19, 20*). The phenotypic markers used to define these subtypes are not mutually exclusive which has complicated the reporting of these cell types by various groups (*21*). Importantly, the defining feature of MDSCs is their ability to suppress immune effector cells, whether they be NK cells or T cells (*21, 22*).

In this study we used multicolour flow cytometry and single-cell RNA sequencing (scRNA-seq) to phenotypically characterize postoperative sx-MDSCs, and using a functional *ex-vivo* co-culture suppression assay, we confirmed these cells suppress NK cell effector functions. Using this assay, we screened a small molecule library to identify molecules that can prevent sx-MDSC suppression. This led us to identify PI3K-γ as the signalling pathway that governs their suppressive machinery and we further demonstrated that targeting this pathway can attenuate postoperative NK cell suppression and cancer metastases.

## RESULTS

### Surgery-induced Myeloid Derived Suppressor Cells in Cancer Patients

The expansion and persistence of immature myeloid cells in the postoperative period has been observed and reported previously (*5, 9, 17, 23*). To corroborate these findings we assessed a cohort of cancer surgery patients (n=55; Table 1) from various cancer types, procedures, sex and ages for changes in immune cell proportions immediately following surgery (postoperative day 1; POD1) using a harmonized multicolour flow cytometry panel for human MDSCs (*19*) (Fig. 1A and B; fig. S1, S2). We focused our analysis on the expansion and characterization of myeloid cells (CD33^+^Lin^-^), which are elevated prior to surgery in cancer patients compared to healthy volunteers and had significantly increased by 1.9-fold on POD1 (Fig. 1C, p<0.0001).

**Figure 1.**
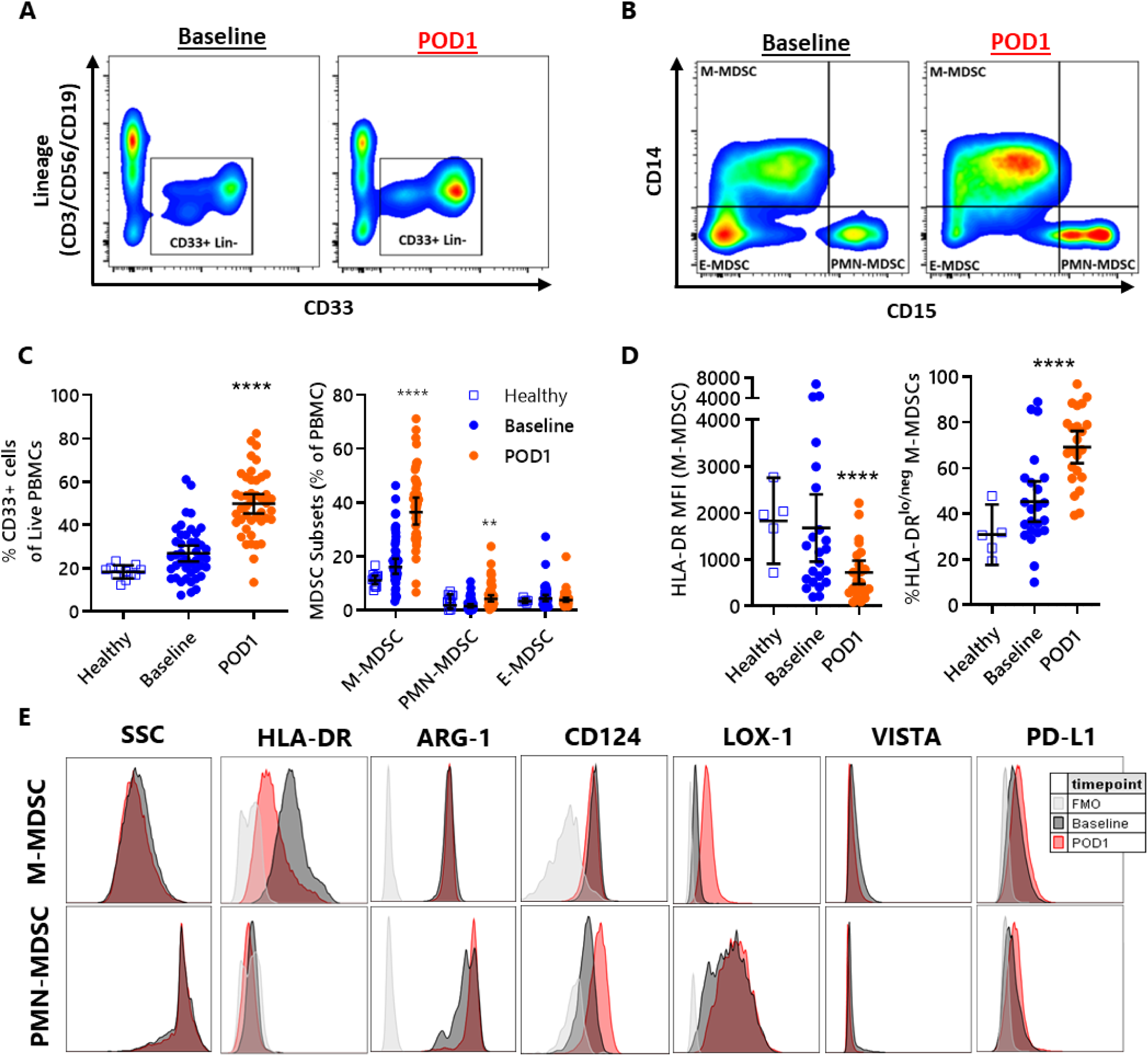
Large expansion of phenotypically defined sx-MDSCs comprised mainly of M-MDSC and PMN-MDSCs immediately after surgery (POD1). **(A)** Representative flow plots showing CD33 vs Lineage (CD3/CD56/CD19) for Baseline (left) and POD1 (right) PBMCs. **(B)** Representative flow plots gated down on CD33^+^Lin^-^ cells showing CD15 vs CD14 expression. **(C)** Healthy controls (n=15), Baseline, and POD1 patients (n=44) proportion of CD33^+^Lin^-^ cells (left) and MDSC subsets (right). **(D)** The MFI of HLA-DR gated on M-MDSCs (left) and the proportion of M-MDSCs that are HLA-DR^lo^ (right). **(E)** Representative histograms of common M-MDSC and PMN-MDSC markers, before and after surgery.

**Table 1.** Patient Demographics for Flow Cytometry.

| Category | Subcategory | Cancer patients (n=55) | Healthy (n=6) |
| --- | --- | --- | --- |
| Sex | Male (n) | 30 | 4 |
|  | Female (n) | 25 | 2 |
| Age - median; range |  | 64; 29-81 | 30; 28-42 |
|  | < 50 years (n) | 8 | 6 |
|  | 50-69 years (n) | 33 | 0 |
|  | > 70 years (n) | 14 | 0 |
| Cancer Type - n | GI | 15 | N/A |
|  | GU | 14 |  |
|  | Prostate* | 12 |  |
|  | Lung | 9 |  |
|  | Sarcoma | 4 |  |
|  | Neuroendocrine | 1 |  |
| Cancer Stage - n (%) | Stage 0 (in situ) | 10 | N/A |
|  | Stage 1 | 14 |  |
|  | Stage 2 | 16 |  |
|  | Stage 3 | 13 |  |
|  | Stage 4 | 2 |  |
| Length of Operation – median; range |  | 3.5; 2-10 | N/A |
| GI – Gastrointestinal (upper and lower GI, & Pancreatic cancer) |  |  |  |
| GU – Genitourinary (kidney, bladder, ureteral, uterine/endometrial) |  |  |  |

In humans, MDSCs are categorized into monocytic (M-MDSC; CD14^+^CD15^lo^HLA-DR^lo^), polymorphonuclear (PMN-MDSC; CD14^-^CD15^hi^HLA-DR^-^), or early stage MDSCs (E-MDSC; CD14^-^CD15^-^) (*24*). In our cancer surgery cohort, the M-MDSC and PMN-MDSC subtypes expanded 2.1-fold and 2.9-fold, respectively, and there was no observed change in the frequency of E-MDSCs after surgery (Fig. 1C). The majority of sx-MDSCs on POD1 were M-MDSCs which accounted for 39.6% (95%CI: 34.8-44.5) of PBMCs, while PMN-MDSCs and E-MDSCs accounted for 6.0% (95%CI: 4.2-8.0) and 4.4% (95%CI: 3.3-5.5), respectively. On POD1, a significant decrease in the MFI of HLA-DR on M-MDSCs was observed, decreasing from an average MFI of 1677 to 724 (p<0.0001), and the proportion of HLA-DR^lo^ M-MDSCs increased significantly from 40% to 66% (p<0.0001) (Fig. 1D, fig. S1).

We also assessed the effect of surgery on MDSC-specific markers such as arginase-1 (Arg1), CD124 (IL-4Rɑ) (*25*), and Lox-1 (*26*), as well as exploratory immune checkpoint markers VISTA (*27*) and PD-L1 (*28*). Although M-MDSCs and PMN-MDSCs have different levels of expression for these markers, they did not significantly increase after surgery (Fig. 1E). Next, we wanted to confirm the suppressive functions of the different sx-MDSC subtypes.

Sx-MDSCs isolated from cancer patients on POD1 (fig. S3) and co-cultured with NK92- MI cells suppressed NK cell cytotoxicity against K562 target tumour cells (Fig. 2A and B). This was not due to NK92 cells targeting the CD33^+^ cells because the viability of CD33^+^ cells was not affected following 6- or 24-hour co-cultures with NK92 alone (Fig. 2B) nor was this due to CD33^+^ cells reducing NK92-MI cell viability (data not shown). Sx- MDSCs on POD1 had greater NK cell suppression compared to MDSCs isolated at Baseline (Fig. 2C). To determine the suppressive capacity of different MDSC subsets, we isolated PMN-MDSCs, M-MDSCs, and also high-density neutrophils (HDNs) on POD1 (Fig. 2D, fig. S3) and co-cultured these cells with NK92 cells. POD1 M-MDSCs suppressed NK cell cytotoxicity to the same extent as bulk-MDSCs, while POD1 PMN-MDSCs only partially suppressed. Notably, HDNs did not result in any suppression (Fig. 2E). Therefore, on POD1, sx-MDSCs are a mixture of M-MDSC (majority) and PMN-MDSC (minority) subsets with different phenotypic features, and surgery-induced M-MDSCs are the primary drivers of NK cell suppression.

**Figure 2.**
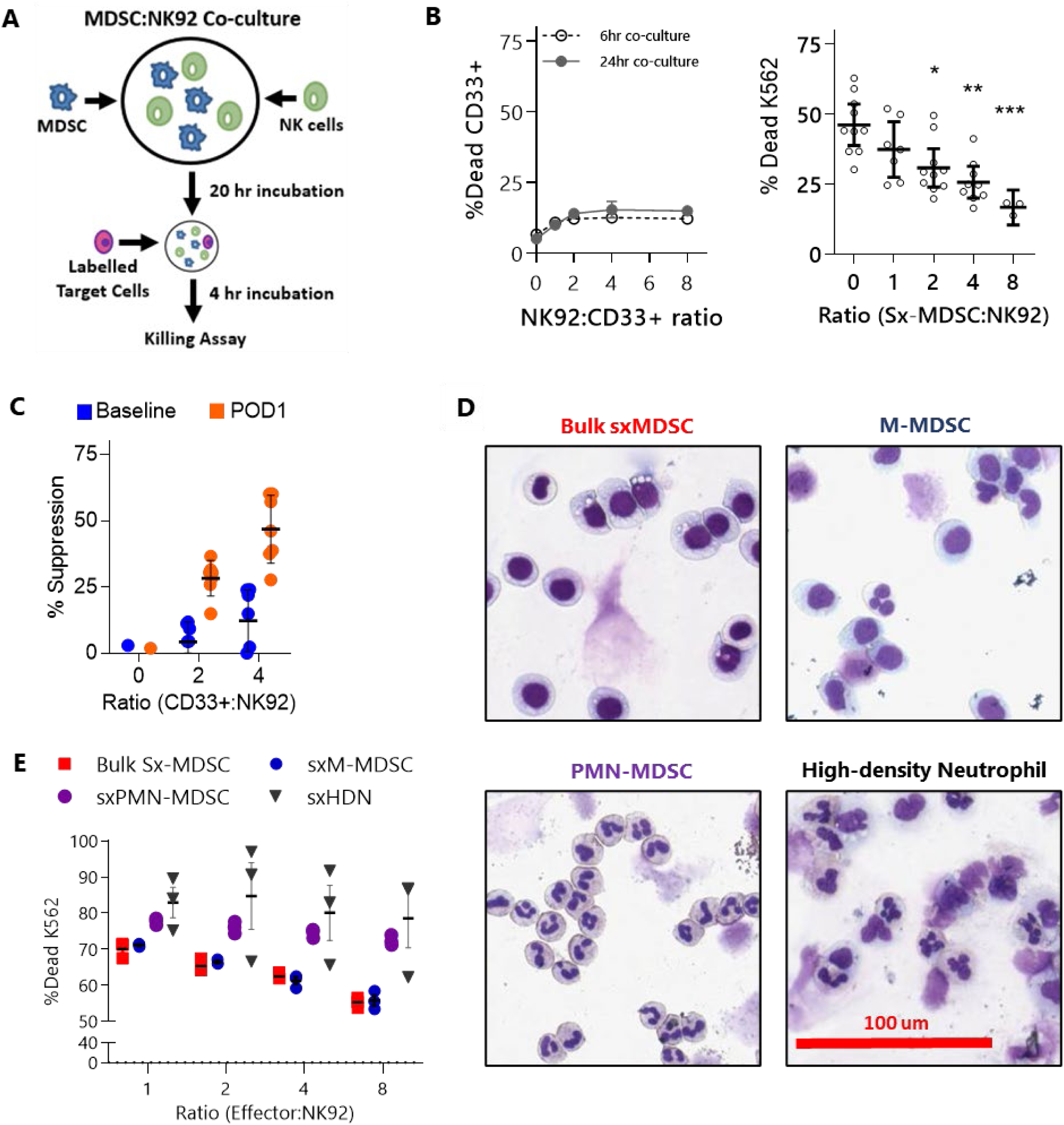
Sx-MDSCs suppress NK92 cytotoxicity. **(A)** Schematic of MDSC-NK92 co-culture suppression assay with CP450-labelled K562 target cells. **(B)** NK92s were co-cultured at increasing ratios with CD33+ cells for 6 or 24 hours (in the absence of K562s) to assess CD33 death by NK92s (left). Sx-MDSCs from POD1 patient samples (n=10) were co-cultured with NK92 cells and NK cytotoxicity was measured as % dead K562 (right). **(C)** % suppression was calculated as the reduction of NK cytotoxicity, normalized to NK cells alone (ratio = 0). CD33+ cells isolated from Baseline and patient-matched POD1 blood were co-cultured with NK92 and K562 targets to measure their suppressive capacity (n=5). **(D)** Representative images of sorted cells stained with Giemsa Wright. **(E)** Bulk Sx-MDSCs, M-MDSCs, PMN-MDSCs, and high-density neutrophils (HDN) from POD1 patients (n=3) co-cultured with NK92 to measure effect on NK cytotoxicity.

### Single-cell RNA sequencing captures significant transcriptional perturbations in monocytes after surgery resembling immunosuppressive MDSCs

To understand the transcriptional changes occurring in the myeloid population after surgery, we performed scRNA-seq on cryopreserved PBMCs from matched baseline and POD1 patient samples (n=6) (Fig. 3A; fig. S4; Table S1). After removing cells with low gene detection (<200 detected genes) and high mitochondrial gene content (>25%), we integrated and clustered the remaining 30,773 cells by underlying cell type (Fig. 3B), resulting in the identification of 10 cell types (Fig. 3C and D). Interestingly, we observed reproducible shifts in the transcriptional profiles of all immune cell types after surgery. Although there were no differences on POD1 in the proportion of NK cells (p=0.8), there was a significant decrease in the %CD3+ cells (0.6 fold-change, p=0.03) and increase in the %CD14+ monocytes (2.0 fold-change, p=0.3) of PBMCs measured by flow cytometry (fig. S4B-D). As the monocytic sx-MDSCs make up the majority of the PBMCs postoperatively (fig. S4D, Fig. 3D and E), we performed differential expression (*29*) followed by gene set enrichment analysis (GSEA) using a collection of query gene sets from the Molecular Signatures Database (GO terms, KEGG pathways, Reactome pathways, and Hallmark gene sets) (*30, 31*) on the monocytic populations. This revealed an upregulation of expected/known responses to surgery such as IL-1 signalling, coagulation, and wound healing (Fig. 3F). Interestingly, there was an increase in the “TNF-ɑ signalling via NF-κB” Hallmark gene set which has been described to be upregulated in immunosuppressive monocytes and MDSCs (*32*). Furthermore, the downregulation of the Reactome pathway “MHC class II antigen presentation” has been described previously (*5*), and supports our flow cytometry observations (Fig. 1D).

**Figure 3.**
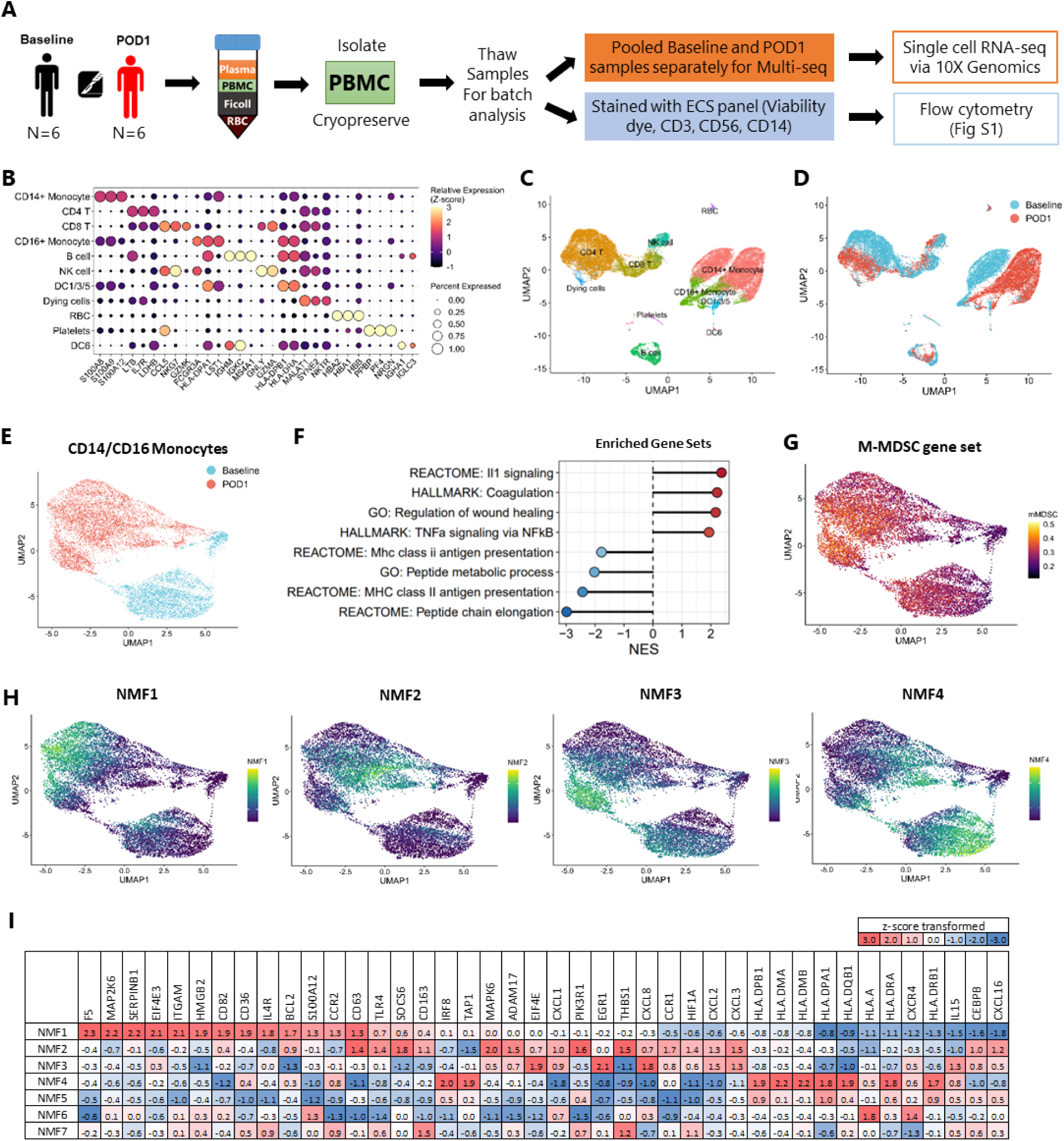
Single cell RNA sequencing (scRNA-seq) of cryopreserved PBMCs before and after surgery reveals drastically altered monocyte/myeloid cell expression profiles on POD1. **(A)** Schematic showing PBMCs (cryopreserved) from 6 matched Baseline and POD1 patients were processed for multiplexed scRNA-seq with the 10x Genomics Chromium platform. **(B)** Dot plot displaying the relative expression of the top 3 marker genes (x-axis) of each cluster (y-axis). **(C)** UMAP plot of scRNA-seq data. Each point corresponds to a single cell and is coloured by cluster. **(D)** Identical UMAP embedding as in (C), with Baseline and POD1 cells labelled. **(E)** UMAP plot of the CD14+ and CD16+ monocyte population at Baseline and POD1. **(F)** Gene set enrichment analysis (GSEA) showing the normalised enrichment scores (NES) of the top upregulated and downregulated pathways. All gene sets are significantly enriched (FDR < 0.05). **(G)** UMAP plot showing activity of an M-MDSC gene set from Alshetaiwai *et al.*; in monocyte populations in Baseline and POD1 samples. **(H)** Non-negative matrix factorization (NMF) plots of gene expression programs for NMF 1, 2, 3 and 4 (see fig. S5 for all NMF programs). **(I)** A heatmap of selected genes driving the various NMF gene expression programs (z-score transformed; ranked according to NMF1).

Although the monocyte clusters separated into two distinct populations (Baseline and POD1, Fig. 3E), we observed that each population retained patterns of phenotypic gradients in their gene expression profiles, suggesting that the monocytes are on a spectrum of differentiation before and after surgery as opposed to being clearly defined postoperative monocytic cell subtypes. This is best exemplified by the observation that similar gradients of gene expression programs can be found in both Baseline and POD1 states. To understand the differences in genes driving a global shift vs. a gradient effect, we used non-negative matrix factorization (NMF) to learn distinct modules of coordinated gene expression (*33*). This resulted in the identification of seven gene expression programs with variable activity in monocytes on Baseline and POD1 (NMF1- 7; fig. S5). Some NMF programs (NMF1, NMF2 and NMF7) were mainly expressed in monocytes on POD1 while other NMF programs (NMF3, NMF4 and NMF6) had a similar expression at Baseline and POD1. Lastly, NMF5 was densely expressed in the CD16+ monocyte populations (fig. S5).

We observed that M-MDSC gene signatures recently identified by scRNA-seq (*34*) were most expressed in the cells using NMF1 and 2 gene expression programs (Fig. 3G and H). Genes contributing to NMF1 and 2 included MDSC-related genes such as *S100a12, Hmgb2, Adam17, Hif1a, Ccr2, Il4r and Serpinb1* (*34–36*). Notably, NMF4 activity was inversely correlated with this signature and driven by expression of genes involved in antigen presentation, including the HLA family genes (e.g. *Hla.dra, Hla.drb1, Hla.dma, Hla.dpa1*) and *Tap1*, corroborating our GSEA results. Lastly, cells highly expressing NMF3 and 7 gene expression programs may be driven by individual patients, ARG12 and ARG19, respectively (fig. S4E and S5A). These NMF programs were also more expressed by POD1 cells and had high expression of genes related to MDSCs (*CEBPB, EIF4E, Trem1*, *Il1b*, *Tnf*, and *Ptgs2*) (*37, 38*). Therefore, our NMF analysis showed that monocytes have varying levels of activity for each gene expression program, but that there was higher activity of NMF1, 2, 3, and 7 on POD1 with similarities to MDSC-related genes (Fig. 3I). These results implicate 4 different activity states of sx-MDSC which each contribute uniquely to the immunosuppressed period after surgery.

### PI3K regulates sx-MDSCs suppressive machinery and is amenable to targeting

To identify pathways regulating the suppressive activity of MDSCs, we performed a series of compound screens using small molecules that affect major biological signalling pathways including PI3K/AKT, TGF-β, VEGF, NFkB, COX, NOS, MMPs, and pathways governing apoptosis and autophagy (adMare Bioinnovations; Fig. 4A). In Screen #1 (Table S2), 147 compounds were incubated in MDSC:NK92 co-cultures for 1 hour at a final concentration of 1μM and the compounds which prevent NK cell suppression by >50% were identified and used in Screen #2 (Table S3). In Screen #2, the compounds that resulted in the greatest reduction (Table S4) in MDSC suppression were compounds that targeted the PI3K pathway, such as LY294002 and AS-252424, suggesting a potential role for PI3K signalling in sx-MDSCs. Numerous reports have shown an important role for PI3K signalling in MDSCs (*39–41*). Therefore, we investigated the role of PI3K in mediating the suppressive activity of sx-MDSC (Fig. 4B and C).

**Figure 4.**
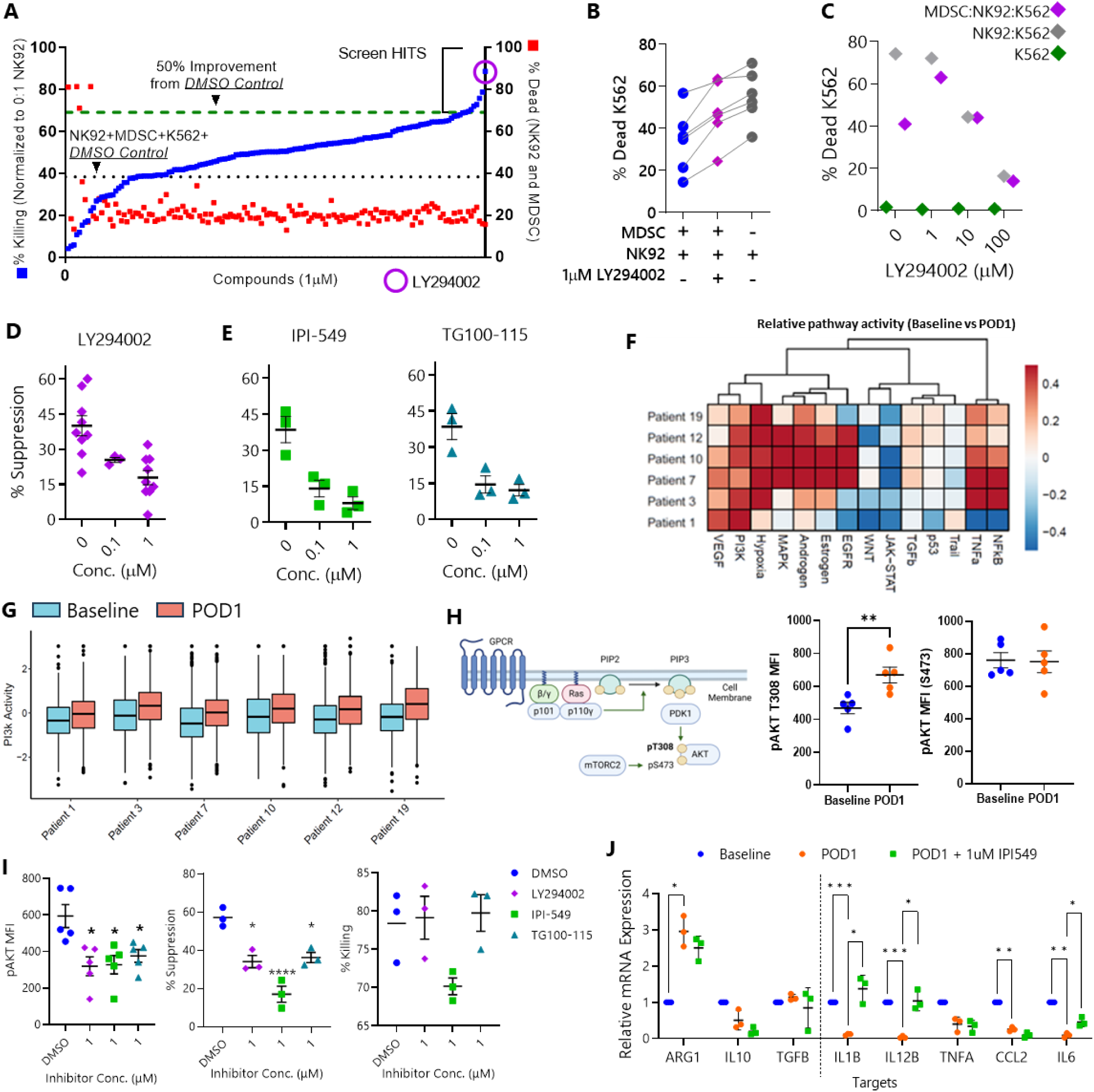
PI3K inhibitors reverse the suppressive effects of sx-MDSCs on NK cells. **(A)** A library of 147 small molecules covering the major cellular signalling pathways were screened (n=4 experiments) at 1μM in the MDSC-NK suppression assay. Compounds that improved NK cell cytotoxicity (blue, left y-axis) by >50% from DMSO control (black dotted line), without impacting NK cell viability (red, right y-axis), were considered hits. LY294002 (purple circle) improved cytotoxicity in all screens and was the top hit in 3/4 screens. **(B)** LY294002 improves NKC (n=6). **(C)** Dose response of LY294002 and the effect on NKC and K562 viability. **(D)** pan-PI3K and **(E)** PI3K-γ specific inhibitors (IPI-549, TG100-115) on MDSC-NK cell suppression. **(F)** Cohort of differentially expressed genes from baseline vs POD1 MDSCs. **(G)** Box plot inference of PI3K pathway activity in baseline vs. POD1 MDSCs. **(H)** Pathway diagram of PI3K signalling and Akt phosphorylation. Phosphorylation status of AKT at the T308 (right) and S473 residue (left) (n=5). **(I)** Effect of PI3K inhibitors on pAKT (T308) MFI (left), MDSC-NK suppression (middle), and NKC (right). **(J)** Effect of IPI-549 on POD1 expression of anti- vs. pro-inflammatory mRNA transcripts, normalized to Baseline. Bar graphs show Mean ± SEM.

Although LY294002 is able to bind to multiple catalytic subunits of class I PI3K heterodimers, the p110-γ subunit is preferentially expressed in myeloid cells (*42*) and is known to regulate immunosuppressive profiles of tumour associated myeloid cells (*39, 41*) . To specifically target PI3K-γ, IPI-549 and TG100-115 (*43*) were used in our co-culture assays. Similar to our findings with LY294002, both of these inhibitors improved NK cell cytotoxicity (Fig. 4D and E) and inhibited Akt phosphorylation in sx-MDSCs (fig. S6A). PI3K-γ specific inhibitors were more effective than LY294002 at reducing phospho-Akt in *ex vivo* whole blood experiments with and without IL-4 stimulation (fig. S6B). Notably, TG100-115 was also able to increase the MFI of HLA-DR when gated on sx-MDSCs in a dose dependent manner, as has been reported by Kaneda et al. (fig. S6B, n=1) (*39*).

### Signalling through the PI3K pathway is increased following surgery

Given the results from the small molecule screen, we used PROGENy to infer relative differences in the signalling pathway activity of monocytic cells in our scRNA-seq. On POD1, there was significant upregulation in several signalling pathways: VEGF, PI3K, hypoxia, MAPK, androgen, NF-κB, and TNF-ɑ. Conversely there was a decrease in JAK-STAT and WNT signalling, while TGF-β, p53 and Trail signalling did not differ significantly after surgery (Fig. 4F). Importantly, PI3K signalling pathway activity was upregulated in all patients (Fig. 4G). Consistent with our functional data, PI3K-regulated transcripts were significantly altered in monocytes after surgery. To determine which NMF program in our monocyte population was associated with elevated PI3K activity, we queried our scRNA-seq against publicly available gene sets of PI3K-regulated genes (*44*). The NMF programs that had the highest correlation with PI3K-regulated genes were NMF1 (*r =* 0.22) and 2 (*r=*0.32), suggesting that these putative sx-MDSC gene programs are also associated with elevated PI3K activity. Therefore, the PI3K signalling pathway is transcriptionally elevated in sx-MDSCs.

Next, we investigated if these transcriptional changes in PI3K were associated with alteration in PI3K signalling. Since PI3K signalling leads to AKT phosphorylation, specifically at residue T308, monitoring the phosphorylation status of this “master regulator” could be a surrogate for PI3K activity (Fig. 4H) (*45*). Phosphorylation of AKT at T308 residue was significantly greater in POD1 M-MDSCs than baseline and there were no changes in S473 phosphorylation. PI3K-γ inhibitor treatment (IPI-549 and TG100-115) led to a dose-dependent reduction in AKT phosphorylation, indicating a reduction in PI3K-γ signalling comparable to LY294002. Moreover, these inhibitors reduced sx-MDSC mediated suppression of NK cell cytotoxicity and did not affect cytotoxicity in the absence of sx-MDSCs (Fig. 4I). However, we note that these inhibitors had a negative effect on cytotoxicity at higher concentrations when plated with NK92 alone (fig. S7).

Finally, we investigated the role of PI3K-γ in mediating the expression of immune effector mRNA transcripts such as anti-inflammatory transcripts, *Arg1*, *Il10*, *Tgfb* as well as pro-inflammatory transcripts*, Il1b*, *Il12b*, *Tnfa, Il6, and Ccl2* in MDSCs isolated from Baseline and POD1 (Fig. 4J). Surgical stress caused a significant decrease in all pro-inflammatory transcripts, *Il1b*, *Il12b, Il6* and *Ccl2* while anti-inflammatory *Arg1* was upregulated. PI3K-γ blockade with IPI-549 treatment in POD1 sx-MDSCs prevented the surgery-induced reduction of pro-inflammatory transcripts *Il1b*, *Il6* and *Il12b*, however it did not rescue *Ccl2* mRNA expression.

### PI3K inhibitors inhibit sx-MDSCs in preclinical murine models of surgical stress

In a pre-clinical model, we have shown that murine PMN-MDSCs (Gr1^+^CD11b^+^Ly6G^+^) and M-MDSCs (Gr1^dim^CD11b^+^Ly6C^+^) increase after surgery (*14*). In the present study, we assessed whether PI3K inhibitors can be used to reduce the suppressive effects of sx-MDSCs in mice. PI3K-γ inhibitors resulted in a dose-dependent reduction in AKT phosphorylation in *ex vivo* murine sx-M-MDSCs and sx-PMN-MDSCs (Fig. 5A). Murine sx-MDSCs stimulated with IL-4 had increased phosphorylation of AKT and served as a positive control. Next, we investigated whether surgical stress alone influenced PI3K-γ pathway activation in murine sx-MDSCs. We observed a significant increase in pAKT MFI in both sx-MDSC subsets on POD1 compared to no surgery controls which was significantly reduced when mice were treated with perioperative TG100-115 (6mg/kg) or IPI-549 (30mg/kg), in M-MDSCs but not PMN-MDSCs (Fig. 5B).

**Figure 5.**
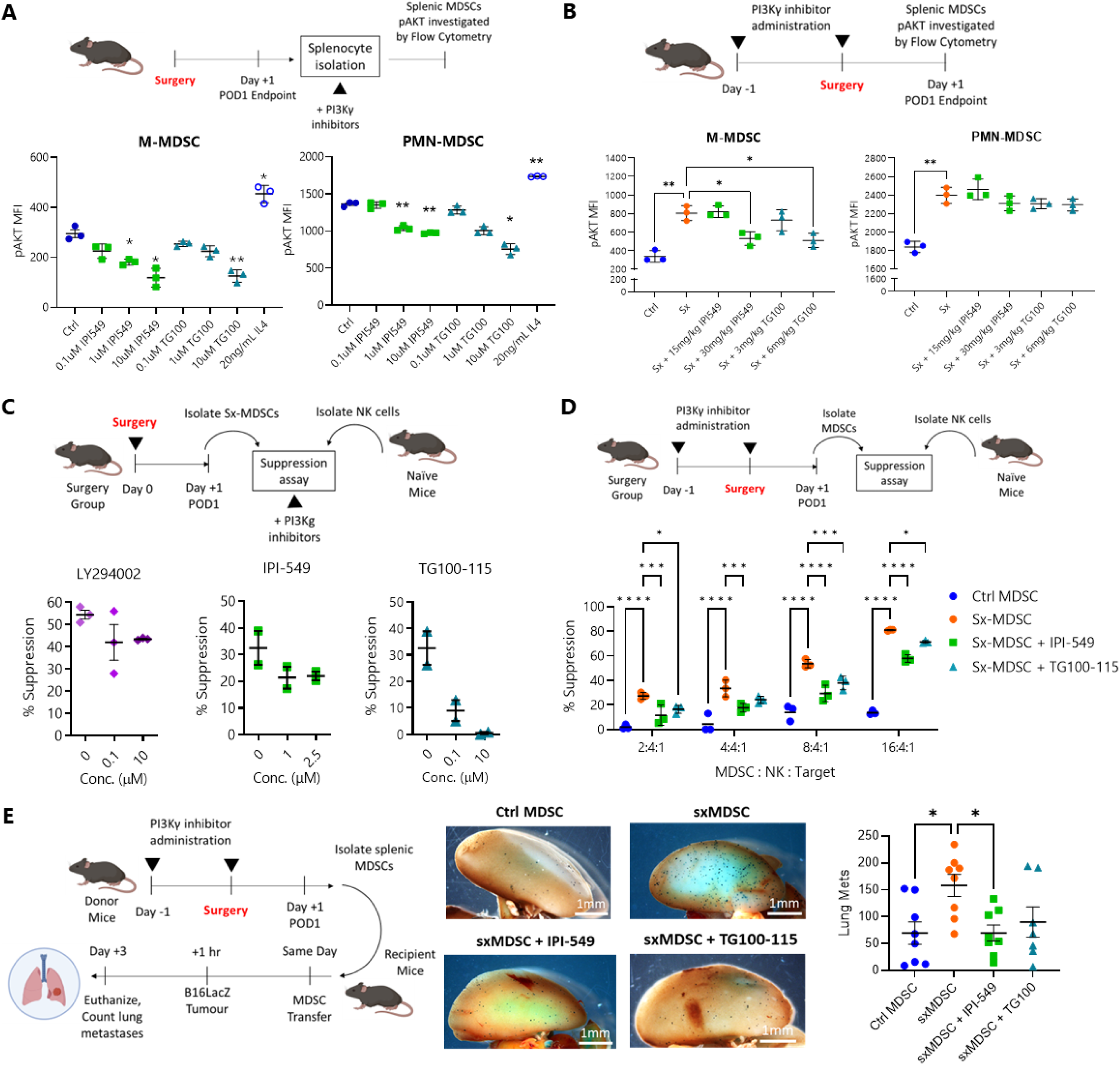
Blockade of PI3K-γ signalling in sx-MDSCs reduces NK cell suppression and metastatic disease in mouse models of surgical stress. **(A)** *Ex vivo* effect of PI3K inhibitors on pAKT (T308) phosphorylation in splenic MDSCs from C57Bl/6 mice (n=4). **(B)** Effect of pre-operative *in vivo* administration of PI3K inhibitors on pAKT (T308) phosphorylation measured on POD1 in splenic MDSCs (n=4). **(C)** Ex vivo effect of PI3K inhibitors on MDSC-NK %suppression (n=3). **(D)** Effect of pre- operative *in vivo* administration of PI3K-γ inhibitors on sx-MDSC suppressive capacity. **(E)** Adoptive transfer of Representative images of lungs are shown.

To assess whether PI3K inhibitors can prevent sx-MDSC mediated suppression of NK cell cytotoxicity, we first tried an *ex-vivo* suppression assay, where MDSCs (Gr1^+^) cells from surgically stressed mice were co-cultured with naïve NK cells from control mice, in the presence of PI3K inhibitors (Fig. 5C). We observed a dose dependent reduction of NK cell cytotoxicity suppression in *ex vivo* co-cultures treated with the PI3K inhibitors. Next, we wanted to know the effect of treating mice perioperatively with TG100-115 or IPI-549. MDSCs from these perioperatively treated mice were then isolated and co-cultured with NK cells from naïve mice (Fig. 5D) in our suppression assay. The sx-MDSCs isolated from mice perioperatively treated with PI3K inhibitors were significantly less suppressive than sx-MDSCs from control mice. However, we observed that systemic perioperative PI3K-γ inhibitor treatment also had a negative effect on NK cell function and led to increased tumour metastases, as well as reduced *ex vivo* NK cell cytotoxicity against Yac-1 targets compared to mice that only received surgery (fig. S8). This effect may be due to off-target effects of PI3K inhibitors on NK cells.

Therefore, in order to assess the effects of PI3K-γ inhibition on sx-MDSCs in isolation, we performed an adoptive transfer of systemically treated sx-MDSCs into naive mice, which were then inoculated with B16F10lacZ tumour cells so that we could measure the resulting lung tumour burden after 3 days (Fig. 5E). The adoptive transfer of sx-MDSCs caused a significant increase in lung metastases compared to no-surgery controls, recapitulating the premetastatic effect of sx-MDSCs, while the adoptive transfer of sx-MDSCs from mice treated with IPI-549 developed significantly fewer lung metastases.

## DISCUSSION

The earliest report of NK cell suppression by a surgery-induced suppressive monocytic cell population was in 1982 by Uchida and colleagues (*46*). Decades later, as our understanding increased, it is likely that Uchida et al. was investigating sx-MDSCs. Only recently has the umbrella term “MDSC” become an accepted nomenclature for the heterogeneous population of immunoregulatory myeloid cells (*47*). Numerous context dependent identifiers have been used to further categorize MDSCs to bring greater clarity and accuracy to the field. This is because factors such as tissue localization, disease state, including cancer subtype and inflammatory conditions, can all contribute to differing MDSC subtypes (*48, 49*).

MDSCs have been best characterized in cancer patients as immature myeloid cells that directly suppress immune effector cells in the tumour microenvironment (*22, 50, 51*). “Surgery-induced” or “trauma-induced” has been used by our group (*17*) and others (*8, 52, 53*) as a prefix for MDSCs that arise from sterile (non-microbial), surgical inflammatory conditions (i.e., inflammatory response to host-derived DAMPs) (*54*). Here, we wanted to determine the contribution of sx-MDSCs, in the inflammatory context of cancer. To that end, we assessed the changes in MDSC populations from a diverse cohort of cancer surgery patients differing in their primary cancer, and surgical procedures (Table 1). This allowed us to assess the *universal* effect of surgery-induced inflammation on postoperative MDSCs. Despite varying starting M-MDSC levels at baseline, the median fold-change was higher in M-MDSCs than PMN-MDSCs across all cancer types, apart from lung cancer surgery patients (fig. S2). In studies where there was a specific cancer type, similar results were reported. Wang et al. (*8*) observed that the M-MDSCs were the main MDSC subtype after surgery in cohort of lung cancer surgery patients. Gao *et al.* (*55*) reported that high systemic inflammation following surgery in hepatocellular carcinoma patients was correlated with a greater expansion of M-MDSCs that was also correlated with a shorter time to recurrence and shorter overall survival. Wiktorin et al. (*53*) found expansion of M-MDSCs in pancreatic cancer surgery patients, linking their presence with a reduced overall survival following surgery as well. In the latter study, PD-L1 cell surface expression increased modestly postoperatively on POD3 to 5 but not on POD1, similar to our results. They also measured a significant increase in ROS on POD1, which we have previously seen in murine sx-MDSC (*17*) but did not measure in human sx-MDSCs in this study.

When comparing PMN-MDSCs and M-MDSCs, PMN-MDSCs had a higher scatter profile compared to M-MDSCs, in addition to greater expression of Arg1, CD124, and Lox1. Lox1 was recently described as a distinguishing biomarker for PMN-MDSCs (*26*) and in our study, Lox1 had a higher postoperative MFI in both sx-MDSC subtypes (Fig. 1E). However, although M-MDSCs also expressed Arg1 and CD124, the MFI of these markers did not increase after surgery in M-MDSCs. M-MDSCs made up the bulk of the sx-MDSC population and had more suppressive capacity when they were separately enriched for, compared to PMN-MDSCs (Fig. 3E).

Single-cell mass cytometry profiling in patients undergoing hip arthroplasty surgery, showed the greatest immune cell perturbations occurred in CD14^+^ monocytes, which resembled the phenotype of immunosuppressive MDSCs (*5*). Similarly, our study showed the most striking differences in transcription were within the monocytic compartment after surgery. We observed that gradients of gene expression patterns were present at baseline and POD1, suggesting that sx-MDSCs are on a continuum of differentiation as opposed to distinct terminal stages of differentiation or maturity (Fig. 3). Querying our data set against a publicly available gene set of M-MDSCs (*34*), we saw that the gene expression profile overlapped with NMF1 and NMF2. The genes that drive these expression programs in our data set were increased DAMPs (e.g. *Hmgb2, S100a8/9/12*) and chemokines (e.g. *Ccr2*) in addition to decreased HLA gene expression (Fig. 3). Interestingly, NMF4 had the inverse gene expression program of NMF 1 and 2, with very high expression of HLA-DR. Since NMF4 activity was mainly expressed in monocytes at baseline (fig. S5B), this supports the hypothesis that after surgery, emergency myelopoiesis results in an expansion of immature myeloid cells that gain a suppressive phenotype because of the postoperative inflammatory context. A limitation of our scRNA-seq experiments is that we are unable to make firm inferences about PMN-MDSCs due to the negative effect of cryopreservation on PBMC viability and recovery of PMN-MDSCs (*19, 56*). Furthermore, specific genes that define MDSC suppressive mechanisms, Arg1 in particular, are untraceable upon freeze-thawing (*51*).

The small molecule screen helped to identify pathways regulating sx-MDSC suppression, an included compounds that targeted TGF-β, VEGF, Raf/MAPK, and NFκB/TNF-ɑ pathways which were transcriptionally upregulated following surgery by scRNA-seq. The compounds GW5074 and Bay11-0785 which inhibit Raf/MAPK and NKκB/TNF-ɑ, respectively, were among the top compounds that inhibited sx-MDSCs (Table S4). The PI3K-γ pathway was pursued because a pan-PI3K inhibitor was the top hit in our screens (Fig. 4A) and was activated in myeloid cells postoperatively based on scRNA-seq (Fig 4G). Furthermore, the PI3K-γ isoform has been reported as the regulator for macrophage polarization and immunoregulatory programs (*39, 40*). Wigcheren et al. (*57*) demonstrated that PI3K-γ inhibition reversed the immature phenotype of MDSCs, shown by increased HLA-DR expression and decreased expression of immunosuppressive markers, MerTK and ROS. De Henau et al. (*40*) demonstrated that selective PI3K-γ isoform inhibition, with IPI-549, reduced the suppressive activity of tumour-associated MDSCs. In addition, IPI-549 overcame resistance to anti-PD-1 as well as anti-CTLA-4 checkpoint blockade therapy, leading to enhanced anti-tumour immunity and tumour regression in mouse models of cancer. Davis et al. (*58*) reported that inhibiting p110γ enhanced the efficacy of anti-PD-L1 immunotherapy in head and neck squamous cell carcinoma by modulating cytotoxic immune activity against tumours and inhibiting MDSC activity. Currently, clinical trials of IPI-549 for triple-negative breast cancer (NCT03719326) and head and neck squamous cell carcinoma (NCT03795610) are ongoing.

We confirmed upregulation of PI3K activity by demonstrating increased phosphorylation of pAKT (T308) in both humans (Fig. 4H) and in our murine model (Fig. 5B), which was reversed with PI3K-γ blockade. Therefore, at the signalling level, PI3K-γ is activated in post-operative M-MDSCs and is amenable to therapeutic blockade. We focused on blockade of the p110-γ catalytic subunit of PI3K because it is the isoform predominantly expressed by myeloid cells and the PI3K-γ specific inhibitors, IPI-549 and TG100-115, were more potent in inhibiting sx-MDSCs *in-vitro* and in-vivo. Similar to prior studies (*39, 40, 58*), we found that sx-MDSCs exhibit a predominantly anti-inflammatory phenotype, characterized by upregulation of Arg1 and downregulation of pro-inflammatory markers such as IL6, IL12B, CCL2, and IL1B. Although PI3K-γ inhibition in sx-MDSCs did not prevent the upregulation of Arg1 expression, it did result in a marked increase of IL6, IL12B, and IL1B (Fig. 4J). This further supports the role of PI3K-γ targeting to prevent or reverse postoperative immune suppression.

This study is the first to demonstrate the therapeutic potential of perioperative PI3K-γ inhibitors in preventing postoperative immune suppression and metastatic recurrence. While systemic delivery of PI3K inhibitors reduced MDSC suppression after surgery, these inhibitors also had negative effects on NK cell activity and resulted increased postoperative tumour burden (fig S8). PI3K-γ plays a critical role in NK cell migration, development, and effector functions(*59, 60*). Moreover, PI3K-γ is required for cytotoxicity, NK cell-target cell interaction and receptor-induced IFNγ production (*59*). Therefore, targeting potent PI3K inhibitors specifically to sx-MDSCs is required to limit off-target effects. Our experiments confirm this because the adoptive transfer of sx-MDSC from mice treated with PI3K-γ inhibitors did not recapitulate the postoperative tumour burden as seen in the no-treatment control group (Fig. 5E). A potential therapeutic strategy to selectively target sx-MDSCs is the use of Antibody Drug Conjugates (ADC), which have conventionally been explored to deliver a cytotoxic payload to tumour cells. In this context, an ADC carrying a PI3K-γ inhibitor could be targeted to unique cell surface proteins on sx-MDSCs, such as CD33/CD14 or Lox1, which is upregulated on sx-MDSCs. In addition, although PI3K-γ has shown significant efficacy in pre-clinical models, little is known about its clinical safety and efficacy. Clinical trials exploring pan-PI3K (*61*) and PI3K-α (*62*), have demonstrated toxicities such as hyperglycaemia, cutaneous reactions, diarrhoea/colitis, pneumonitis, (*63*). Therefore, targeted sx-MDSC-specific approaches may also mitigate toxicities associated with systemic PI3K-γ blockade.

In summary, the perioperative period is a dynamic landscape occupied by immunosuppressive monocytic sx-MDSCs which have heterogeneous transcriptional profiles. Scoring gene sets from MDSCs generated in different inflammatory settings will help identify the context specific attributes of MDSCs and bring greater clarity and accuracy to the field. Currently, no perioperative therapy is approved to target sx-MDSCs, but we have shown preliminary evidence that sx-MDSCs rely on PI3K signalling for their suppressive activity. Indeed, PI3K controls the transcriptional regulation of the immunosuppressive programming in sx-MDSCs, however, given the critical role of PI3K signalling in anti-tumour immunity, particularly in NK cell function, strategies to specifically target sx-MDSCs need to be explored. Moreover, given the rapid activation of sx-MDSCs soon after surgery with resultant NK cell dysfunction, pre-operative PI3K blockade in sx-MDSCs may pre-emptively prevent this immunosuppressive phenotype from arising. One may question how a relatively short period of profound immune suppression could lead to the discovery of metastases months-to-years-later, but recent studies have clearly demonstrated that the detection of circulating tumour DNA within 3-6 weeks after surgical resection for cancer is highly predictive of recurrence in colorectal (*64*), lung (*65*), breast (*66*), and pancreatic cancer (*67*). Supporting a functional immune system to clear microscopic disease in the early postoperative period is a highly promising strategy to reduce recurrences, providing clinical rationale for a therapeutic strategy that directly targets the underlying mechanism behind postoperative immune suppression.

## MATERIALS AND METHODS

### Study Design

To characterize sx-MDSCs, we used peripheral blood mononuclear cells (PBMCs) or “ACK” (Ammonium-Chloride-Potassium) lysed whole blood from patients (n=55) enrolled under the Perioperative Human Blood and Tissue Specimen Collection Program (PHBSP; OHSN-REB# 2011884-01H). Inclusion and exclusion criteria were determined prospectively. Consenting participants over the age of 18 undergoing a surgical procedure for their cancer treatment were included in the study. Patients who were previously treated with chemotherapy, radiation, or immunotherapy, and received blood transfusions during surgery were excluded from the study. The patient samples collected under this study protocol were used in all experiments, excluding the scRNA-seq (Table 1).

To control for inter-assay variability, antibodies were titrated and stained following a standard flow cytometry staining protocol using previously determined gates and voltages. Voltages for excitation/emission spectra were routinely assessed to ensure that MFI values could be compared between baseline and post-operative day (POD) 1, and across different patient samples on different days of experimentation. Lastly, a core panel of antibodies were used to phenotype MDSCs by flow cytometry, with the capacity to include additional exploratory markers. Patient samples were used in multiple experiments to ensure the ethical and efficient use of patient materials.

To explore the transcriptional changes occurring after surgery in sx-MDSCs, we performed scRNA-seq from six cryopreserved PBMCs from abdominal cancer surgery patients (Supplemental Table 1) enrolled in the PERIOP-02 clinical trial (NCT02987296; OHSN-REB# 20160732-01H). These patients were instructed to take a supplement enriched in arginine (n=3) or an isocaloric/isonitrogenous control supplement (n=3) TID for 5 days as part of the trial (*68*). To account for the potential impact of the differences in treatments, we performed a separate analysis comparing the transcriptional changes between groups and did not find any effect attributable to the supplement given (fig. S4). Furthermore, the patient samples selected for the scRNA-seq had similar demographic, procedural and surgical outcome data (Supplemental Table 1). Therefore, we combined all six patients in our final analysis to increase our statistical power.

### Human Blood Sample Collection and Processing

Patient (Baseline and POD1) and healthy volunteer peripheral blood samples (20-40mL) were drawn at The Ottawa Hospital with informed consent under the following clinical protocols: *i)* OHSN-REB# 20160732-01H and *ii)* OHSN-REB# 2011884-01H. Blood was drawn into BD Vacutainer Sodium-Heparin coated tubes and processed within 2 hours by Ficoll density centrifugation (GE Healthcare #17-1440-03).

### Antibodies for Phenotyping by Flow Cytometry

The panel of MDSC markers were chosen based on published guidelines to harmonize human MDSC reporting led by the Association for Cancer Immunotherapy, Cancer Immunoguiding Program (*19*) -with slight modifications. Freshly isolated PBMCs were first labelled with a fixable viability stain (BV510; BD #564406) in PBS at room temperature for 10 minutes. Next, an extracellular MDSC antibody mastermix was added to the PBMCs and simultaneously stained with the viability dye for an additional 20 minutes at 4℃. The antibodies in the MDSC mastermix were used at individually titrated dilutions and included: CD33 clone P67.6 (Pe-Cy7, Biolegend #366617), CD14 clone MθP9 (APC-Cy7, BD #561709), CD11b clone ICRF44 (AF700, Novus #FAB1699N), HLA-DR clone L243 (APC, Biolegend #307609), CD15 clone MMA (efluor450, eBioscience #48-0158-41), CD124 clone 25463 (PE, R&D systems #FAB230P-025), and lineage markers CD3 clone UCHT1 (FITC, Biolegend #300440), CD56 clone NCAM16.2 (FITC, BD Biosciences #340410) and CD19 clone HIB19 (FITC, Biolegend #302205) (*19*). Exploratory extracellular markers included LOX-1 clone 15C4 (PE, Biolegend #358603), VISTA clone 730804 (AF700, R&D systems #FAB71261N), and PD-L1 clone MIH1 (PE, eBioscience #12-5983-42). Following viability and extracellular staining, PBMCs were washed and resuspended in 1% PFA and acquired by flow cytometry within 48 hours. For intracellular staining, the BD Cytofix/Cytoperm™ kit (BD #554714) was used after viability and extracellular staining and cells were incubated in Perm/Wash buffer with Arginase-1 (APC, R&D systems #IC5868A) for 30 minutes at 4℃. Samples were acquired on the BD Fortessa LSRII and analyzed with FlowJo v10.

### Cell Lines and Isolation of sx-MDSC Subsets and High-density Neutrophils

PBMCs from whole blood taken on POD1 were processed through two workflows to isolate M-MDSCs (CD33^+^CD14^+^CD15^-^) and PMN-MDSCs (CD33^dim^CD14^-/dim^CD15^+^) (fig. S3). For M-MDSC isolation, CD33^+^ cells were first separated using CD33^+^ microbeads (Miltenyi #130-045-501) via immunomagnetic bead separation. Similarly, PMN-MDSCs were isolated by first obtaining CD15^+^ cells using CD15^+^ microbeads (Miltenyi #130-046-601). Both starting populations were then stained with a simplified immunophenotyping panel containing: CD33 clone P67.6 (Pe-Cy7, Biolegend #366617), CD14 clone MθP9 (APC-Cy7, BD #561709), CD15 clone W6D3 (PE, BD Biosciences #562371), L/D (APC, Thermo Scientific #L34975). The stained CD33^+^ and CD15^+^ cells underwent fluorescence activated cell sorting (FACS) using the Sony MA900 Multi-Application Cell Sorter to isolate live M-MDSCs (CD33^+^CD14^+^CD15^-^) and PMN-MDSCs (CD33^dim^CD14^-/dim^CD15^+^), respectively.

High density neutrophils (HDNs) were isolated via double density centrifugation by overlaying 6mL whole blood on top of 3mL Histopaque 1077 (Sigma-Aldrich #10771), which was first overlaid on top of 3mL Histopaque 1119 (Sigma-Aldrich #11191). After centrifugation at 700 x G for 30 minutes with brakes off, the PBMCs were collected from the upper interface, followed by HDN collection from the bottom interface.

### MDSC:NK92 Suppression Assay

K562 and NK92-MI cell lines were purchased from ATCC and maintained in complete RPMI (RPMI+10% FBS; “cRPMI”). MDSC mediated NK cell suppression was measured by isolating MDSCs from surgery patients on POD1 and incubating them in cRPMI at increasing ratios with the IL-2 independent NK cell line, NK92-MI for 20 hours at 37℃. Following incubation, K562 target cells labelled with Cell Proliferation Dye efluor 450 (CP450, eBioscience #65-0842-90) were added to the MDSC:NK co-cultures for 4 hours. NK cell cytotoxicity was then measured by adding Ethidium homodimer (EtHD, Invitrogen™ #E1169) to each well just prior to acquiring the samples by flow cytometry. NK cell cytotoxicity was reported as %dead K562 (EtHD+ CP450+). To calculate %MDSC suppression, we used the following equation:

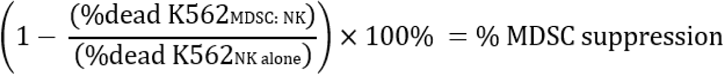

### Giemsa-Wright Staining and Microscopy

Freshly isolated PBMCs were washed twice in MilliQ H_2_O and resuspended in 20μL of MilliQ H_2_O. 5-10μL of PBMCs were pipetted onto a glass slide and gently smeared with the long edge of a 20μL pipette tip. Slides were fixed with 100% MeOH and air dried for 2 minutes. At this point, 0.5 to 1mL of undiluted stock Giemsa-Wright stain (Sigma #WG80-2.5L) was pipetted onto the slides and left for 2 minutes to stain. Afterwards, excess Giemsa-Wright stain was poured off and the slides were submerged into a diluted Giemsa-Wright stain (1:20, stain:MilliQ H_2_O) for 5 minutes. Slides were then gently submerged into MilliQ H_2_O to rinse and left to air dry. Slides were viewed and imaged the same day (Nikon Te2000e, The Ottawa Hospital Research Institute).

### Single-cell RNA Sequencing Library Preparation and Sequencing

Cryopreserved and matched Baseline/POD1 PBMC patient samples (n=6) were used for single-cell RNA-sequencing (scRNA-seq) on the 10x Genomics Chromium Single Cell 3’ platform (StemCore Laboratories, the Ottawa Hospital Research Institute). Baseline and POD1 samples were multiplexed into separate pools using MULTI-seq before being processed on two lanes of the Chromium platform superloaded to target a 20,000 cell yield per lane. Briefly, each sample was labelled for 10 minutes with 200nM of a unique DNA barcode hybridized to complementary lipid-modified DNA enabling anchoring to the cells’ plasma membrane. Each sample was washed twice with PBS+1% BSA prior to pooling. Gene expression libraries were prepared following the standard 10x Genomics protocol and separate MULTI-seq barcode libraries were isolated by size selection and PCR amplified as described by McGinnis et al. (https://github.com/chris-mcginnis-ucsf/MULTI-seq). Libraries were sequenced using NextSeq500 (Illumina) high-output 75-cycle runs. Gene expression libraries were sequenced to a depth of approximately 20,000 reads/cell and MULTI-seq barcode libraries were sequenced to a depth of approximately 1000 reads per cell, which is sufficient for sample annotation.

### scRNA-seq Data Processing and Pipeline

Following sequencing, fastq files were generated using the Cell Ranger mkfastq tool (10x Genomics). Gene expression libraries were processed using the Cell Ranger count function, aligning to the human transcriptome (GRCh38) with default parameters other than explicitly setting --expect-cells 20000. MULTI-seq barcode fastq files were processed using the deMULTIplex tool developed for MULTI-seq (https://github.com/chris-mcginnis-ucsf/MULTI-seq), providing a sample annotation for each cell barcode.

Gene expression libraries were imported into R and processed with Seurat (*69*). Cells containing high proportions of mitochondrial reads were first removed before splitting the data into separate 12 Seurat objects representing each unique sample. Each sample was normalized using SCTransform prior to integration using Seurat’s SCTransform integration pipeline (*69, 70*). The integrated data was then processed with principal component analysis (PCA) and clustered using the Louvain algorithm (resolution=0.15) on a neighbour graph derived from the first 30 principal components. Deriving cluster identities based on the integrated data ensured that clusters were not associated with variability of each patient or experimental groupings (ie. baseline/POD1). Each cluster was annotated based on expression of canonical markers of PBMC populations.

Following clustering, unintegrated data was re-processed using SCTransform, PCA, and UMAP embedding, providing a low-dimensional representation of the data capturing biological variability of interest while retaining cluster annotations that represent the underlying cell type. We then used the R tool muscat to perform differential state analysis between POD1 and baseline samples within each cluster (*29*).

### Signalling Pathway Activity Inference

The R package PROGENy was used to infer the activity of 14 signalling pathways in each cell of the data (*71*). PROGENy derives activity scores using a regression model trained on data from transcriptional profiles from hundreds of signalling perturbation experiments. After inferring activity scores, we tested for pathways with differential activity between cells from POD1 and baseline samples using a simple linear model.

### Gene Set Enrichment Analysis

The R package fgsea was used to perform gene set enrichment analysis on genes ranked by their average log fold change from the differential state analysis for individual cell types (*72*). Lists of all GO terms, KEGG pathways, Reactome pathways, and Hallmark gene sets were acquired from the Molecular Signatures Database (MSigDB) (*30, 31*) http://paperpile.com/b/rMuHDH/vW8em. All gene sets presented in the manuscript are significantly enriched (adjusted p-value < 0.05) and normalized enrichment scores (NES) are presented.

### Identification of Coordinated Gene Expression Programs

To identify coordinated gene expression programs that represent continuous phenotypic gradients, we applied consensus non-negative matrix factorization (NMF) to normalized gene expression counts for the top 2000 variable genes (based on variance computed by SCTransform), as implemented in the NMF tool described by Kotliar et al. (*33*). To identify the appropriate number of programs to identify (factors; k), we used NMF to perform 100 iterations of NMF for each k from 2-10. k=7 provided stable factorizations with low error and was used for downstream analysis (fig. S5). Consensus NMF then identifies a consensus factorization based on the 100 iterations for a given k. The “cell usage” and coefficient matrices resulting from the factorization were then imported into R. As a gene’s coefficient is ultimately dependent on its average expression level, we Z-score-transformed each gene’s coefficients across the NMF programs and ranked genes by their Z-score value for each NMF program. To gain insight into which biological phenotypes may be associated with each NMF program, we calculated correlated cells’ NMF program usages with gene set scores from MSigDB and publications of interest.

### Gene Set Scoring and Autocorrelation

Gene set scoring was performed using Vision (*73*). Similar to complementary tools, Vision computes a score for query gene sets based on the average expression of each gene relative to background control gene sets. Vision also calculates an autocorrelation score (1-Geary’s C) for each gene set, providing insight into whether gene set activity represents coordinated variation in latent space (kNN graph from 50-dimensional PCA space) or if the scores are randomly distributed.

### Small Molecule Screen for MDSC Antagonists

Our MDSC antagonists screens were separated into 2 screens (Screen #1 and Screen #2) which were each performed twice. Screen #1 included 147 unique small molecule compounds plated in singlets at 1µM (Supplemental Table 2). Screen #1 was created in collaboration with adMare Bioinnovations (British Columbia, Canada). The library was plated in 96 v-bottom plates and shipped on dry ice via priority courier to the OHRI where the suppression assays were done immediately upon receiving the compounds. Each plate contained their own DMSO and media only wells in which control samples were plated. Sx-MDSCs were isolated from cancer surgery patients on POD1 and plated at a 4:1 sx-MDSC:NK92 cell ratio +/- drug compounds for 24 hours prior to adding CP450-labelled K562 cells. Compounds which led to the greatest improvement in NK cell cytotoxicity (hits) from Screen #1 informed the creation of Screen #2 which included a refined list of only 40 compounds, plated in duplicates, at 1µM and 10µM (Supplemental Table 3). Drugs were plated in a randomized order and kept blinded from experimenters until after completing the analysis.

### Phospho-Signalling Assay

For *in vitro* phospho-AKT staining, PBMCs (humans) or splenocytes (mouse) were incubated for 3 hours with or without PI3K-γ inhibitor treatment in cRPMI at 37°C in a 5% CO_2_ chamber. During the last 15 minutes of incubation, MDSCs were stained with an antibody panel for immunophenotyping. Splenocytes underwent a simultaneous Lyse/Fix step using Lyse/Fix buffer (BD Biosciences # 558049) while PBMCs were fixed with fixation buffer (BD Biosciences #554655), and both were incubated for 10 minutes at 37°C. Samples were then permeabilized according to the BD Phosflow protocol III (*74*) using Perm buffer III (BD Biosciences #558050). Cells were washed twice with flow buffer and then stained with AKT pT308 Clone D25E6 (PE, Cell Signalling Technology #13038S) and AKT pS473 Clone M89-61 (AF647, BD Biosciences #561670) for 1 hour on ice in the dark. A final wash was performed with flow buffer and the phosphorylation status of AKT was acquired on the BD Fortessa LSRII and analysed with FlowJo v10.

### Reverse Transcriptase (RT) Quantitative PCR (qPCR)

MDSCs were diluted to a concentration of 1×106 cells/mL in the presence or absence of 1μM IPI-549 (Selleckchem #S8330) for 24 hours at 37°C. 2×10^6^ MDSCs were used for RNA extraction using the Rnaeasy Plus Kit (Qiagen # 4134), according to the manufacturer’s protocol. Extracted RNA was quantified on the ThermoScientific NanoDrop One Spectrophotometer. cDNA was prepared using 1μg RNA with the Applied Biosystems™ High-Capacity cDNA Reverse Transcription Kit with RNase Inhibitor (Thermo Fisher Scientific #4374966), according to the manufacturer’s protocol. PCR primers were titrated, and optimal melting temperature was determined by preliminary melt curve experiment. Sybr green-based qPCR was performed using human primers to *Arg1, Ccl2, Il10, Il6, Tgfb, Il1b, Il12b, Tbp* and *Tnfa*, using the SsoAdvanced™ Universal SYBR® Green Supermix (Bio-Rad #1725271). mRNA levels were normalized to *Tbp* (ΔCt = Ct^gene of interest^ – Ct^Tbp^). mRNA expression was normalized to baseline samples (ΔΔCt = 2^-(ΔCt^sample^ – ΔCt^baseline^)).

### Murine Studies

All C57Bl/6 mice were purchased from Charles River Laboratories (Wilmington, VA) and housed under strict pathogen-free conditions at the University of Ottawa Animal Care and Veterinary Services (ACVS) facility (Ottawa, ON). Murine protocols complied with the Canadian Council on Animal Care guidelines and were approved by the University of Ottawa Animal Research Ethics Board prior to initiating experiments.

*In vivo,* formulations for TG100-115 were prepared at 3mg/kg or 6mg/kg of body weight in PBS with 30% (v/v) PEG-300 (Sigma-Aldrich #8.07484). IPI-549 was prepared at 15mg/kg or 30mg/kg of body weight in PBS with 40% (v/v) PEG-400 (Sigma-Aldrich #25322-68-3). Formulations were prepared fresh, and mice were treated daily from 1 day before surgery (Day −1) up to POD1. IPI-549 was administered by oral gavage twice daily while TG100-115 was administered once daily intraperitoneally. 200μL of either formulation was administered.

### Murine Surgical Stress Model

Murine surgical stress was induced by performing a laparotomy and invasive left nephrectomy followed by abdominal closure using 5-0 Polysorb suture (Covidien #SL5687G) and skin staples as previously described (*14, 17*). On POD1 or POD3, the mice were sacrificed by lethal buprenorphine IP injection at 0.1 mg/kg body weight, followed by cervical dislocation. Spleens were harvested via the left upper quadrant abdominal incision and splenocytes were dissociated through sterile 70 µm Cell Strainers (Thermo Fisher Scientific #08-771-2). Splenocytes were washed once in cRPMI, counted, and used for subsequent experiments.

### Murine MDSC Suppression Assay

Sx-MDSCs were isolated using the MDSC Isolation Kit (Miltenyi #130-094-538) from surgically stressed mice on POD1. NK cells from no surgery control mice were isolated using the EasySep^TM^ NK Isolation Kit (Stemcell #100-0960). NK cells and Sx-MDSCs were co-cultured for overnight in the presence of 100U/mL of recombinant IL-2 (Thermo Fisher Scientific #CB-40043). YAC-1 cells were stained with CP450 and used as targets for the NK cells.

### Assessment of Postoperative Metastases

B16F10-LacZ melanoma cells (3×10^5^ cells, minimum 90% viability) were injected via intravenous (IV) tail vein into C57Bl/6 mice. Three days following the tumour challenge, our standardized LacZ staining protocol was used for lung metastases (*14*). On the first day, lungs were washed twice in Wash Buffer (1M magnesium chloride, 1% deoxycholate, 2% nonidet-P40, 0.1M sodium phosphate buffer pH 7.3), and then stained overnight at 37°C with X-Gal^TM^ (Thermo Fisher Scientific #R0404) solution, 25 mg/mL in Dimethylsulfoxide. The next day, lungs were washed twice with Wash Buffer and incubated for 24 hours at 4°C. On the third day, lungs were transferred into 10% buffered formalin for permanent fixation. Lungs were imaged using the Axiovision software v4.8 (Zeiss) on the Zeiss SteREO^TM^ Discovery Modular Microscope (Zeiss). From these electronic images, lung metastases were quantified using Fiji ImageJ.

### MDSC Adoptive Transfer

To assess the effects of MDSC-specific inhibition of PI3K-γ, we utilised an MDSC adoptive transfer method (*17*). In this study, MDSC-donor mice received TG100-115, IPI-549, or no treatment (n = 8 per group), one day after surgery. Additionally, there was a MDSC-Donor group that did not receive treatment or surgery. On POD1, MDSCs were isolated and resuspended to 1×10^7^ cells/mL concentration. Next, 5×10^6^ isolated MDSCs were adoptively transferred to each MDSC-recipient mice from their respective MDSC- donor experimental group, except for the no-transfer control group. One hour after the adoptive MDSC transfer, MDSC-recipient mice were tumour challenged with B16F10-LacZ, and lung metastases were counted on day 3.

### Statistical Analysis

Statistical analysis was performed in GraphPad Prism 8. Unpaired, nonparametric Mann-Whitney U test was used when comparing two groups and paired Wilcoxon rank-sum test was used to compare two matched samples (i.e., baseline vs POD1). Multiple comparisons were tested with nonparametric Kruskal-Wallis tests. P values were considered significant when p < 0.05. Bonferroni corrections were applied when multiple queries were applied to a data set.

## List of Supplementary Materials

**Supplemental Table 1.** Patient data for samples used in scRNA-Seq.

**Supplemental Table 2.** Screen #1 compound list.

**Supplemental Table 3.** Screen #2 compound list.

**Supplemental Table 4.** List of top hits from Screens #1A, 1B, 2A, and 2B.

**Supplemental Figure S1.** Gating strategy for MDSC immunophenotyping.

**Supplemental Figure S2.** Subgroup analysis on expansion of M-MDSCs.

**Supplemental Figure S3.** Workflow for isolating sx-MDSC subtypes.

**Supplemental Figure S4.** scRNA-seq extended data 1.

**Supplemental Figure S5.** scRNA-seq extended data 2 – NMF plots.

**Supplemental Figure S6.** Phospho-flow cytometry for pAKT and pS6 gated on sx-MDSCs +/- PI3K inhibitors.

**Supplemental Figure S7.** Validation of top candidate compounds following.

**Supplemental Figure S8.** Systemic delivery of PI3K-γ inhibitors does not prevent metastases in our B16F10LacZ murine model of surgical stress.

## Acknowledgements

We would like to thank J.N., A.J., and M.S., for enrolling and scheduling patients into this study. We thank E.R. for helping in optimizing the Giemsa Wright staining protocol. Special thanks to I.S., A.V.R., J.L., P.B., T.P., and J.Y., for their collaboration throughout the project.

## Author contributions

Conceptualization: RCA, MAK, LA, GT

Methodology: LA, GT, DC, AM, MM, CTS, EC, MAK, RCA

Investigation: RCA, LA, GT

Funding acquisition: RCA

Supervision: RCA, MAK

Writing – original draft: LA, GT, DC, RCA

Writing – review & editing: NK, MA, BV, MAK, RCA

## Data and materials availability

All data are available in the main text of the supplementary materials. Any data, code, and materials presented in this study may be made available to researchers by request to the corresponding author.

