## Supplemental Tables and Figures for "Preventing surgery induced immune suppression and metastases by inhibiting PI3K-gamma signalling in Myeloid-Derived Suppressor Cells"

### SUPPLEMENTARY TABLES AND FIGURES

#### Supplemental Table 1. Patient data for samples used in scRNA-Seq

| Patient ID | Sex | Age | Stage | CCI | Trial Arm *** | Type of surgery | OR Pathology details | NKC (POD1/BL) | SX-MDSCs (POD1/BL) |
|---|---|---|---|---|---|---|---|---|---|
| ARG01 | M | 56 | II | 2 | A | Lap | Adenocarcinoma | 0.93 | 2.46 |
| ARG03 | M | 66 | I | 4 | A | Lap | Adenocarcinoma | 0.36 | 1.95 |
| ARG19 | M | 66 | II | 4 | A | Lap | Adenocarcinoma | N/A | 2.23 |
| ARG07 | M | 65 | II | 4 | B | Open | Adenocarcinoma | 0.67 | 1.36 |
| ARG10 | M | 57 | I | 3 | B | Lap | Invasive carcinoma in ascending polyp | 0.18 | 2.46 |
| ARG12 | M | 79 | II | 5 | B | Lap | Adenocarcinoma | 0.40 | 1.29 |

*Footnote text
**Data taken from the PERIOP-02 Clinical Trial Registered (NCT02987296)
***Participants enrolled were randomized to receive either an arginine enriched supplement ("A") or isocaloric/isonitrogenous control supplement ("B") for 5 days prior to surgery.

### Supplemental Table 2. Screen #1 compound list

Supplemental Table 2. Screen #1 compound list

| Compounds #1-75 | Chemical name | Targeted Pathway |
|---|---|---|
| 1 | CGP 57380 | MAPK inhibitor |
| 2 | Tyrphostin A9 | PDGF |
| 3 | I-OMe-Tyrphostin AG 538 | IGF1R Inhibitor |
| 4 | AC-55649 | RARβ2 receptor agonist |
| 5 | U-73122 | Phospholipase C inhibitor |
| 6 | Imiquimod | TLR7 agonist |
| 7 | SQ 22536 | Adenylyl Cyclase inhibitor |
| 8 | Tyrphostin AG 808 | tyrosine kinase inhibitor |
| 9 | Pifithrin-mu | Bcl-xL and Bcl-2 |
| 10 | Tyrphostin AG 490 | Jak-2 protein tyrosine kinase inhibitor |
| 11 | DMSO | Control |
| 12 | SU 4312 | Neuronal NOS inhibitor |
| 13 | SCH-202676 hydrobromide | GPCR ligand inhibitor |
| 14 | Sulindac | NSAID |
| 15 | Retinoic acid | enhances cell maturation |
| 16 | Rottlerin | Protein kinase Cδ (PKCδ) inhibitor. |
| 17 | Stattic | STAT3 inhibitor |
| 18 | A3 hydrochloride | Non-selective casein kinase inhibitor. |
| 19 | SMER28 | enhancer of rapamycin |
| 20 | Bay 11-7082 | inhibitor of IκBα phosphorylation |
| 21 | Quercetin dihydrate | antioxidant flavonoid |
| 22 | ARP 101 | MMP-2 inhibitor |
| 23 | SB 216763 | GSK-3 inhibitor |
| 24 | Piceatannol | promotes glucose uptake |
| 25 | Olomoucine | CDK inhibitor |
| 26 | Nimesulide | NSAID |
| 27 | NG-Nitro-L-arginine methyl ester hydrochloride | iNOS inhibitor |
| 28 | GW9662 | PPARγ antagonist |
| 29 | Myricetin | Flavonoid antioxidant |
| 30 | Meclofenamic acid sodium | COX inhibitor |
| 31 | beta-Lapachone | Modulates NAD metabolism |
| 32 | SB-525334 | TGFβ receptor I inhibitor |
| 33 | LY-294,002 hydrochloride | pan-PI3K inhibitor |
| 34 | SD-169 | MAPK inhibitor |
| 35 | 1-(5-Isoquinolinylsulfonyl)-2-methylpiperazine dihydrochloride | PKC inhibitor |
| 36 | Olvanil | synthetic analogue of capsaicin |
| 37 | LFM-A13 | BTK inhibitor |
| 38 | 3-Isobutyl-1-methylxanthine | cAMP and cGMP inhibitor |
| 39 | ML-7 | Myosin light chain kinase inhibitor |
| 40 | Imazodan | phoshodiesterase III inhibitor |
| 41 | DMSO | Control |
| 42 | JFD00244 | SIRT2 inhibitor |
| 43 | Ibudilast | PDE4 inhibitor |
| 44 | MNS | tyrosine kinase inhibitor |
| 45 | HA-1004 hydrochloride | inhibitor of protein kinase G (PKG) and PKA |
| 46 | HA-100 | PKA, PKC, and PKG inhibitor |
| 47 | Emodin | tyrosine kinase inhibitor |
| 48 | JX401 | p38 MAP Kinase Inhibitor VI |
| 49 | GW7647 | PPARα agonist |
| 50 | Genipin | Prevents NO production |
| 51 | GW5074 | cRaf1 kinase inhibitor |
| 52 | GW2974 | EGFR / ErbB-2 inhibitor |
| 53 | AS 604850 | ATP-competitive PI3Kγ inhibitor |
| 54 | GW1929 | PPAR-γ activator |
| 55 | S-Ethylisothiourea hydrobromide | iNOS, eNOS, and nNOS inhibitors |
| 56 | Forskolin | Protein kinase A agonist |
| 57 | SP600125 | JNK2 inhibitor |
| 58 | 7-Cyclopentyl-5-(4-phenoxy)phenyl-7H-pyrrolo[2,3-d]pyrimidin-4-ylamine | Lck inhibitor |
| 59 | Diclofenac sodium | NSAID |
| 60 | Cytidine 5'-diphosphocholine sodium salt hydrate | synthesis of phosphatidylcholine |
| 61 | 3,7-Dimethyl-1-propargylxanthine | A2 adenosine receptor antagonist |
| 62 | Imperatorin | acetylcholinesterase inhibitor |
| 63 | Enoximone | PDE3 Inhibitor |
| 64 | PD 169316 | p38 MAP kinase inhibitor |
| 65 | Clodronic acid | Calcium modulator |
| 66 | SANT-1 | Inhibits Sonic hedgehog signalling |
| 67 | 1,7-Dimethylxanthine | Adenosine receptor ligand |
| 68 | Cambinol | inhibits NAD-dependant deacetylases |
| 69 | Capsazepine | TRPV1 receptor antagonist |
| 70 | Diacylglycerol Kinase Inhibitor II | Diacylglycerol Kinase Inhibitor II |
| 71 | Y-27632 dihydrochloride | Rho kinase inhibitor |
| 72 | Daphnetin | EGFR , PKA, and PKC inhibtor |
| 73 | CGS-15943 | adenosine receptors A1 and A2A antagonist |
| 74 | 2-Chloroadenosine | adenosine receptor agonist. |
| 75 | Cilostamide | PDE3 Inhibitor |

Supplemental Table 2: Screen #1 compound list (Continued)

| Compounds #76-150 | Chemical name | Targeted Pathway |
|---|---|---|
| 76 | 9-cyclopentyladenine | adenylyl cyclase inhibitor |
| 77 | Chelerythrine chloride | inhibitor of protein kinase C |
| 78 | BRL 50481 | PDE7 Inhibitor |
| 79 | ML-9 | Myosin light chain kinase inhibitor |
| 80 | BWB70C | 5-lipoxygenase inhibitor |
| 81 | L-Cycloserine | serine palmitoyltransferase inhibitor |
| 82 | BTO-1 | polo-like kinase |
| 83 | CP55940 | non-selective cannabinoid receptor agonist |
| 84 | Bay 11-7085 | inhibitor of IκBα phosphorylation |
| 85 | SB 202190 | 5-lipoxygenase inhibitor |
| 86 | AS-252424 | Pi3K-gamma inhibitor |
| 87 | Bromoenol lactone | inhibitor of calcium-independent phospholipase A2 |
| 88 | TBBz | Casein Kinase-2 (CK2) inhibitor |
| 89 | Diacylglycerol kinase inhibitor I | Diacylglycerol kinase inhibitor I |
| 90 | DMSO | Control |
| 91 | 2-(2-Aminoethyl)isothiourea dihydrobromide | iNOS inhibitor |
| 92 | N-arachidonylglycine | GPR18 agonist |
| 93 | Tryptamine hydrochloride | antagonizes 5-HT serotonin receptors |
| 94 | Aminoguanidine hemisulfate | iNOS inhibitor |
| 95 | Acetylsalicylic acid | COX1/2 inhibitor |
| 96 | 5-azacytidine | cytidine analog |
| 97 | YM 976 | PDE4 inhibitor |
| 98 | DL-alpha-Methyl-p-tyrosine | tyrosine hydroxylase enzyme inhibitor |
| 99 | 6-Methoxy-1,2,3,4-tetrahydro-9H-pyrido[3,4b] indole | indole |
| 100 | Acetamide | anti-microbial, antifungal |
| 101 | Amantadine hydrochloride | nicotinic antagonist |
| 102 | GABA | neurotransmitter |
| 103 | Gabaculine hydrochloride | analog of GABA |
| 104 | O-(Carboxymethyl)hydroxylamine hemihydrochloride | forms oximes |
| 105 | (±)-2-Amino-7-phosphonoheptanoic acid | NMDA glutamate receptor antagonist |
| 106 | N-Acetylprocainamide hydrochloride | Class III antiarrhythmic |
| 107 | Actinonin | MMP inhibitor |
| 108 | S(-)-p-Bromotetramisole oxalate | inhibitor of alkaline phosphatase |
| 109 | TMB-8 hydrochloride | antagonist of nicotinic acetylcholine receptors |
| 110 | L-azetidine-2-carboxylic acid | inhibitor of collagen synthesis |
| 111 | S-(p-Azidophenacyl)glutathione | inhibit glyoxalase and glutathione S-transferase |
| 112 | Acetyl-beta-methylcholine chloride | Muscarinic acetylcholine receptor antagonist |
| 113 | AA-861 | 5-Lipoxygenase inhibitor |
| 114 | Azathioprine | prodrug of 6-mercaptopurine |
| 115 | L-732,138 | Neurokinin-1 receptor antagonist |
| 116 | Amifostine | free radical scavenger and detoxifier |
| 117 | Atropine methyl bromide | muscarinic receptor (mAChR) antagonist |
| 118 | Azelaic acid | tyrosinase inhibitor |
| 119 | Atropine methyl nitrate | Muscarinic acetylcholine receptor antagonist |
| 120 | (±)-Norepinephrine (+)bitartrate | alpha- and beta-adrenergic receptors |
| 121 | Aurintricarboxylic acid | inhibitor of ribonuclease and topoisomerase II |
| 122 | 3-Aminopropionitrile fumarate | lysyl oxidase (LOX) inhibitor |
| 123 | 1-Aminobenzotriazole | inhibitor of cytochrome P450 and chloroperoxidase |
| 124 | Sandoz 58-035 | Acyl-CoA:cholesterol acyltransferase (ACAT) inhibitor |
| 125 | Acetylthiocholine chloride | nicotinic acetylcholine receptor agonist |
| 126 | A-315456 | α1D-adrenoceptor antagonist |
| 127 | Agmatine sulfate | Nitric oxide modulator, NADPH oxidase activator |
| 128 | Arcaine sulfate | NMDA glutamate receptor antagonist |
| 129 | 4-Amino-1,8-naphthalimide | PARP inhibitor |
| 130 | (±)-2-Amino-4-phosphonobutyric acid | agonist for the group III metabotropic glutamate receptors |
| 131 | Apigenin | Flavonoid, inhibits MAPK, ERK, JNK and p38 |
| 132 | 3-Amino-1-propanesulfonic acid sodium | GABAA receptor agonist |
| 133 | (±)-2-Amino-3-phosphonopropionic acid | Metabotropic glutamate receptor antagonist |
| 134 | 4-Androsten-4-ol-3,17-dione | unknown |
| 135 | GR 46611 | 5-HT1D agonist |
| 136 | 4-Aminobenzamidine dihydrochloride | fibrinogen receptor antagonists |
| 137 | (±)-Nipecotic acid | inhibitor of uptake of γ-aminobutyric acid (GABA) |
| 138 | Atropine sulfate | antagonist of muscarinic receptors |
| 139 | 3-aminobenzamide | PARP inhibitor |
| 140 | N-Acetyl-5-hydroxytryptamine | precursor of melatonin |
| 141 | 5-(N-Ethyl-N-isopropyl)amiloride | Sodium-hydrogen exchanger inhibitor |
| 142 | 10058-F4 | c-Myc inhibitor |
| 143 | Amiprilose hydrochloride | Inhibits PLA2 substrate availability |
| 144 | 5-(N-Methyl-N-isobutyl)amiloride | nhibitor of Na+/H+ antiporter |
| 145 | Arecoline hydrobromide | agonist of the muscarinic acetylcholine receptors |
| 146 | N-Phenylanthranilic acid | NSAID |
| 147 | S-(4-Nitrobenzyl)-6-thioguanosine | Potent adenosine uptake inhibitor |
| 148 | TLR7 Agonist | TLR7 Agonist |
| 149 | ABT-737 | inhibits Bcl-2 and Bcl-xL |
| 150 | AT406 | IAP Inhibitor |

**Supplemental Table 3. Screen #2 compound list.**

Supplemental Table 3. Screen #2 compound list.

| Compounds #1-40 | Chemical name | Targeted Pathway | Compound # from Screen 1 |
|---|---|---|---|
| 1 | Salubrinal | inhibitor of eIF2α phosphatase | N/A |
| 2 | Luteolin | antioxidant flavonoid | N/A |
| 3 | Ebselen | glutathione peroxidase mimetic | N/A |
| 4 | 4BPA | ER stress inhibitor | N/A |
| 5 | LY294002 | pan-PI3K inhibitor | 33 |
| 6 | LCK Inhibitor | Lck inhibitor | N/A |
| 7 | ABT-737 | inhibits Bcl-2 and Bcl-xL | 149 |
| 8 | AT406 | IAP Inhibitor | 150 |
| 9 | Tyrphostin A9 | PDGF | 2 |
| 10 | U-73122 | PLC inhibitor | 5 |
| 11 | DMSO | Control | 11 |
| 12 | ARP 101 | MMP-2 inhibitor | 22 |
| 13 | Quercetin dihydrate | antioxidant flavonoid | 21 |
| 14 | Imazodan | phoshodiesterase III inhibitor | 40 |
| 15 | MNS | tyrosine kinase inhibitor | 44 |
| 16 | HA-1004 hydrochloride | inhibitor of protein kinase G (PKG) and PKA | 45 |
| 17 | GW5074 | cRaf1 kinase inhibitor | 51 |
| 18 | GW2974 | EGFR / ErbB-2 inhibitor | 52 |
| 19 | AS 604850 | ATP-competitive PI3Kγ inhibitor | 53 |
| 20 | GW1929 | PPAR-γ activator | 54 |
| 21 | Clodronic acid | Calcium modulator, used to deplete BM macrophages | 65 |
| 22 | SANT-1 | Inhibits Sonic hedgehog signalling | 66 |
| 23 | 1,7-Dimethylxanthine | Adenosine receptor ligand | 67 |
| 24 | Cambinol | inhibits NAD-dependant deacetylases | 68 |
| 25 | L-Cycloserine | serine palmitoyltransferase inhibitor | 81 |
| 26 | BTO-1 | polo-like kinase | 82 |
| 27 | CP55940 | non-selective cannabinoid receptor agonist | 83 |
| 28 | Bay 11-7085 | inhibitor of IκBα phosphorylation | 84 |
| 29 | AS-252424 | PI3K-gamma inhibitor | 86 |
| 30 | Bromoenol lactone | inhibitor of calcium-independent phospholipase A2 | 87 |
| 31 | TBBz | Casein Kinase-2 (CK2) inhibitor | 88 |
| 32 | Diacylglycerol kinase inhibitor I | Diacylglycerol kinase inhibitor I | 89 |
| 33 | Aurintricarboxylic acid | inhibitor of ribonuclease and topoisomerase II | 121 |
| 34 | 3-Aminopropionitrile fumarate | lysyl oxidase (LOX) inhibitor | 122 |
| 35 | 1-Aminobenzotriazole | inhibitor of cytochrome P450 and chloroperoxidase | 123 |
| 36 | Sandoz 58-035 | Acyl-CoA:cholesterol acyltransferase (ACAT) inhibitor | 124 |
| 37 | Acetylthiocholine chloride | nicotinic acetylcholine receptor agonist | 125 |
| 38 | A-315456 | α1D-adrenoceptor antagonist | 126 |
| 39 | Agmatine sulfate | Nitric oxide modulator, NADPH oxidase activator | 127 |
| 40 | Arcaine sulfate | NMDA glutamate receptor antagonist | 128 |

### Supplemental Table 4. List of top hits from Screens #1A, 1B, 2A, and 2B.

Supplemental Table 4. List of top hits from Screens #1A, 1B, 2A, and 2B

Screen #1 (Supplemental Table 2)

| Screen | Compound # Screen 1 | Compound | Target | Avg % Dead K562 | Dead normalized | % suppression | >50% improved? |
|---|---|---|---|---|---|---|---|
| S1A | Controls | NK92:K562 (max killing) | | 44.3 | 100% | 0% | |
| S1A | Controls | NK92:MDSC:K562 (MDSC suppression) | | 17.0 | 38% | 62% | Cutoff = 31% |
| S1A | 33 | LY-294,002 hydrochloride | pan-PI3K inhibitor | 39.3 | 89% | 11% | yes |
| S1A | 58 | 7-Cyclopentyl-5-(4-phenoxy)phenyl-7H-pyrrolo[2,3-d]pyrimidin-4-ylamine | Lck inhibitor | 34.9 | 79% | 21% | yes |
| S1A | 52 | GW2974 | EGFR / ErbB-2 inhibitor | 33.6 | 76% | 24% | yes |
| S1A | 143 | Amiprilose hydrochloride | Inhibits PLA2 substrate availability | 32.3 | 73% | 27% | yes |
| S1A | 51 | GW5074 | cRaf1 kinase inhibitor | 31.9 | 72% | 28% | yes |
| S1A | 132 | 3-Amino-1-propanesulfonic acid sodium | GABAA receptor agonist | 31.2 | 70% | 30% | yes |
| S1A | 69 | Capsazepine | TRPV1 receptor antagonist | 30.8 | 69% | 31% | yes |
| S1B | Controls | NK92:K562 (max killing) | | 53.7 | 100% | 0% | |
| S1B | Controls | NK92:MDSC:K562 (MDSC suppression) | | 28.1 | 52% | 48% | Cutoff = 24% |
| S1B | 20 | Bay 11-7082 | inhibitor of IκBα phosphorylation | 46.6 | 87% | 13% | yes |
| S1B | 139 | 3-aminobenzamide | PARP inhibitor | 45.9 | 85% | 15% | yes |
| S1B | 71 | Y-27632 dihydrochloride | Rho kinase inhibitor | 44.3 | 83% | 17% | yes |
| S1B | 74 | 2-Chloroadenosine | adenosine receptor agonist | 41.7 | 78% | 22% | yes |
| S1B | 133 | (±)-2-Amino-3-phosphonopropionic acid | metabotropic glutamate receptor agonist | 41.3 | 77% | 23% | yes |
| S1B | 141 | 5-(N-Ethyl-N-isopropyl)amiloride | Sodium-hydrogen exchanger inhibitor | 40.9 | 76% | 24% | no |
| S1B | 33 | LY-294,002 hydrochloride | pan-PI3K inhibitor | 39.6 | 74% | 26% | no |

Screen #2 (Supplemental Table 3)

| Screen | Compound # Screen 2 | Compound | Target | Avg % Dead K562 | Dead normalized | % suppression | >50% improvement? |
|---|---|---|---|---|---|---|---|
| S2A | Controls | NK92:K562 (max killing) | | 56.7 | 100% | 0% | |
| S2A | Controls | NK92:MDSC:K562 (MDSC suppression) | | 29.8 | 53% | 47% | Cutoff = 23.5% |
| S2A | 17 | GW5074 | cRaf1 kinase inhibitor | 44.9 | 79% | 21% | yes |
| S2A | 5 | LY294002 | pan-PI3K inhibitor | 44.7 | 79% | 21% | yes |
| S2A | 10 | U-73122 | PLC inhibitor | 44.1 | 78% | 22% | yes |
| *S2A* | *28* | *Bay 11-7085* | *inhibitor of IκBα phosphorylation* | *42.5* | *75%* | *25%* | no |
| *S2A* | *4* | *4BPA* | *ER stress inhibitor* | *42.0* | *74%* | *26%* | no |
| *S2A* | *21* | *Clodronic acid* | *Calcium modulator, depletes BM macrophages* | *41.3* | *73%* | *27%* | no |
| S2B | Controls | NK92:K562 (max killing) | | 35 | 100% | 0% | |
| S2B | Controls | NK92:MDSC:K562 (MDSC suppression) | | 11.5 | 33% | 67% | Cutoff = 33.5% |
| S2B | 5 | LY294002 | pan-PI3K inhibitor | 26.9 | 77% | 23% | yes |
| S2B | 29 | AS-252424 | PI3K-gamma inhibitor | 25.1 | 72% | 28% | yes |
| S2B | 22 | SANT-1 | Inhibits Sonic hedgehog signalling | 23.3 | 67% | 33% | yes |
| *S2B* | *21* | *Clodronic acid* | *Calcium modulator, depletes BM macrophages* | *23.2* | *66%* | *34%* | no |
| *S2B* | *28* | *Bay 11-7085* | *inhibitor of IκBα phosphorylation* | *23.2* | *66%* | *34%* | no |
| *S2B* | *10* | *U-73122* | *PLC inhibitor* | *23.0* | *66%* | *34%* | no |

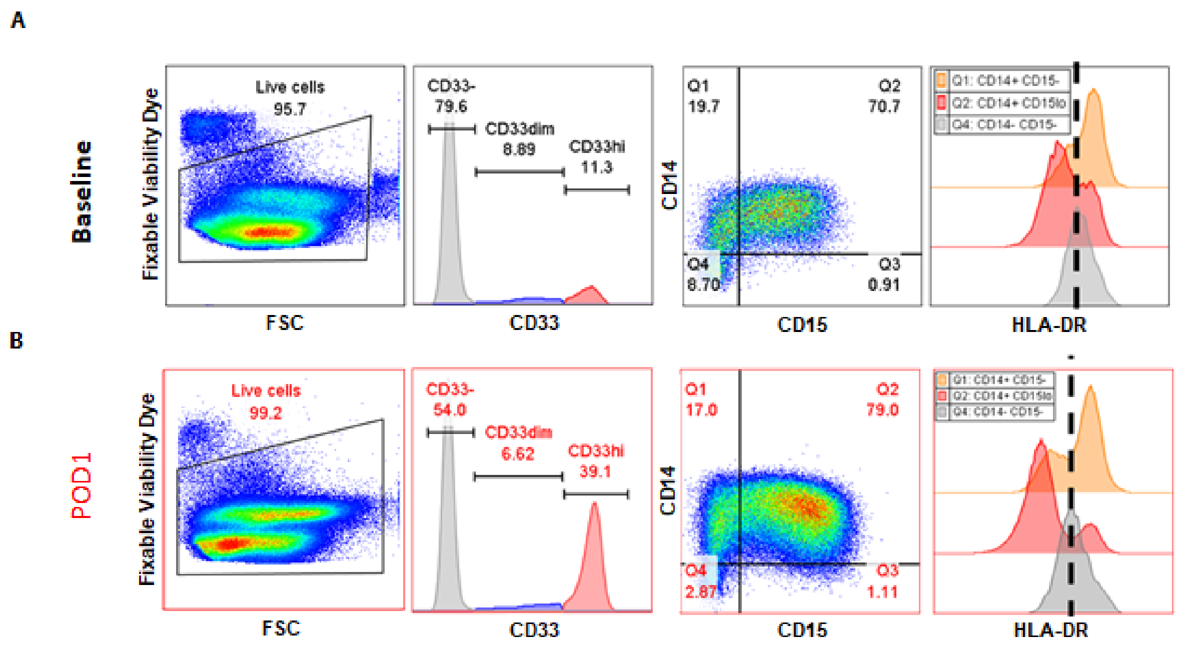


**Supplemental Figure S1. Gating strategy for MDSC immunophenotyping.** Patient PBMCs (baseline or POD1) or murine splenocytes (no surgery or surgery) were stained for phenotypic characterization by flow cytometry immediately after isolation. **(A-B)** Doublets, debris, and dead cells were excluded and live cells were analysed. CD33$^{hi}$ cells were gated on and then analysed for CD14 vs CD15 and HLA-DR expression. The HLA-DR$^{lo}$ cutoff is shown as a black dotted line.

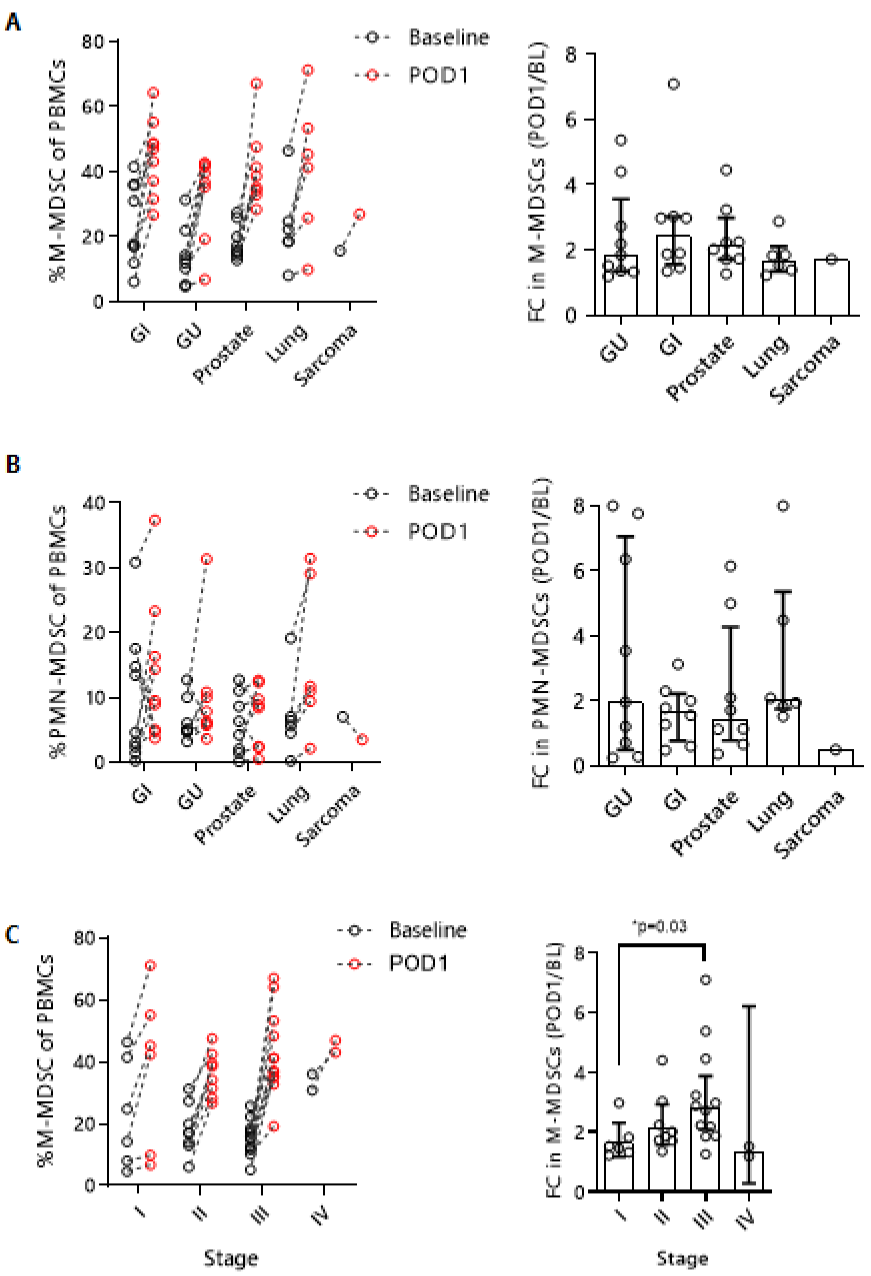


**Supplemental Figure S2. Subgroup analysis on expansion of M-MDSCs.**
**(A)** The effect of various cancer surgery on the %M-MDSCs (left, CD33$^{+}$Lin-CD14$^{+}$CD15$^{lo}$HLA-DR$^{lo}$) and the fold change on POD1 (right). **(B)** %PMN-MDSCs (left, CD33$^{+}$Lin$^{-}$CD14$^{-}$CD15$^{hi}$) and the fold change on POD1 (right). **(C)** Stratifying by %M-MDSCs (left) and the FC in M-MDSC according to cancer stage.

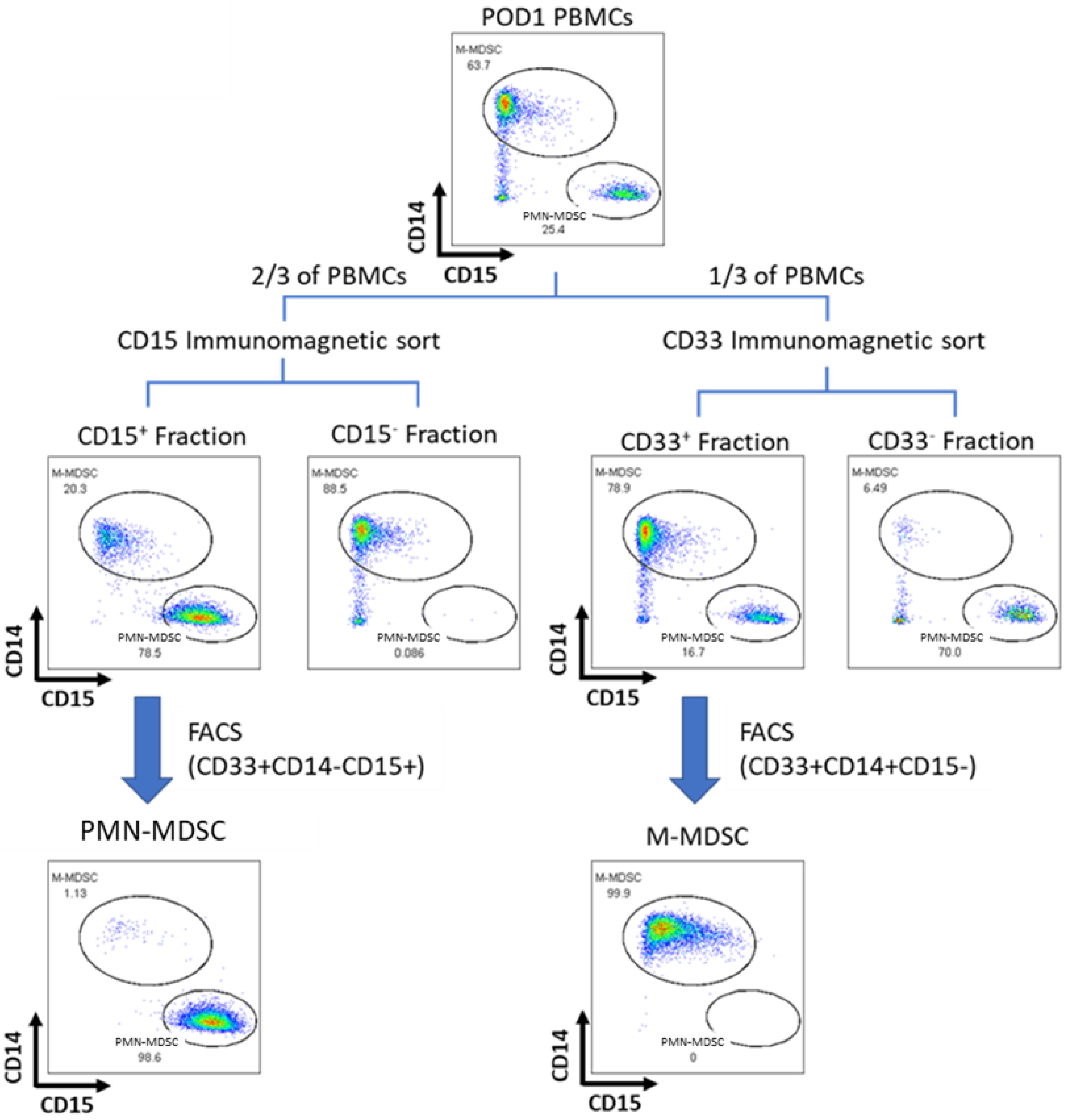


**Supplemental Figure S3. Workflow for isolating sx-MDSC subtypes.**
Gating strategy for FACS of MDSC subpopulations. Cell debris, doublets, dead, and CD33- cells were gated out. CD14+CD15- (M-MDSC) and CD14-CD15+ (G-MDSCS) were gated within the CD33+ cell gate. CD14 vs CD15 flow plot from M- and G-MDSC FACS strategy.

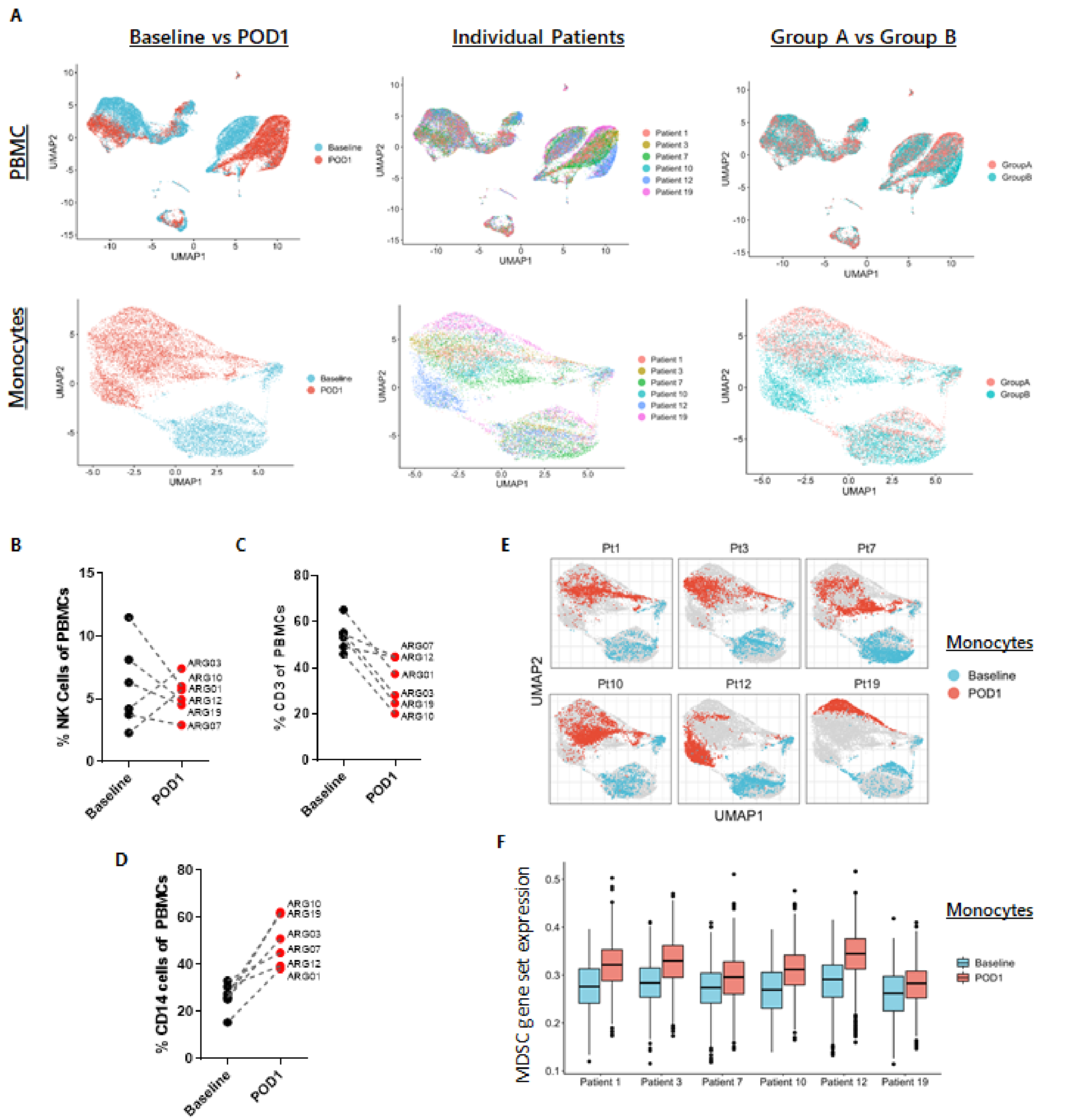


**Supplemental Figure S4. scRNA-seq extended data 1.**
**(A)** UMAP plots of all PBMCs (upper) or monocyte (bottom) populations grouped by: Baseline vs POD1 (left); individual patient (middle); or Group A vs Group B (right). **(B-D)** PBMC samples used for scRNA-seq were simultaneously phenotyped by flow cytometry for changes in the proportion of **(B)** CD3-CD56+ NK cells, **(C)** CD3 T cells, and **(D)** CD14 monocytes before and after surgery. **(E)** Overlaying the contribution of each individual patient set as grouped by Baseline vs POD1 in the PBMC UMAP plots. **(F)** Comparing the expression of MDSC gene set scores for individual patient monocyte populations on Baseline and POD1 (greater = more MDSC-like).

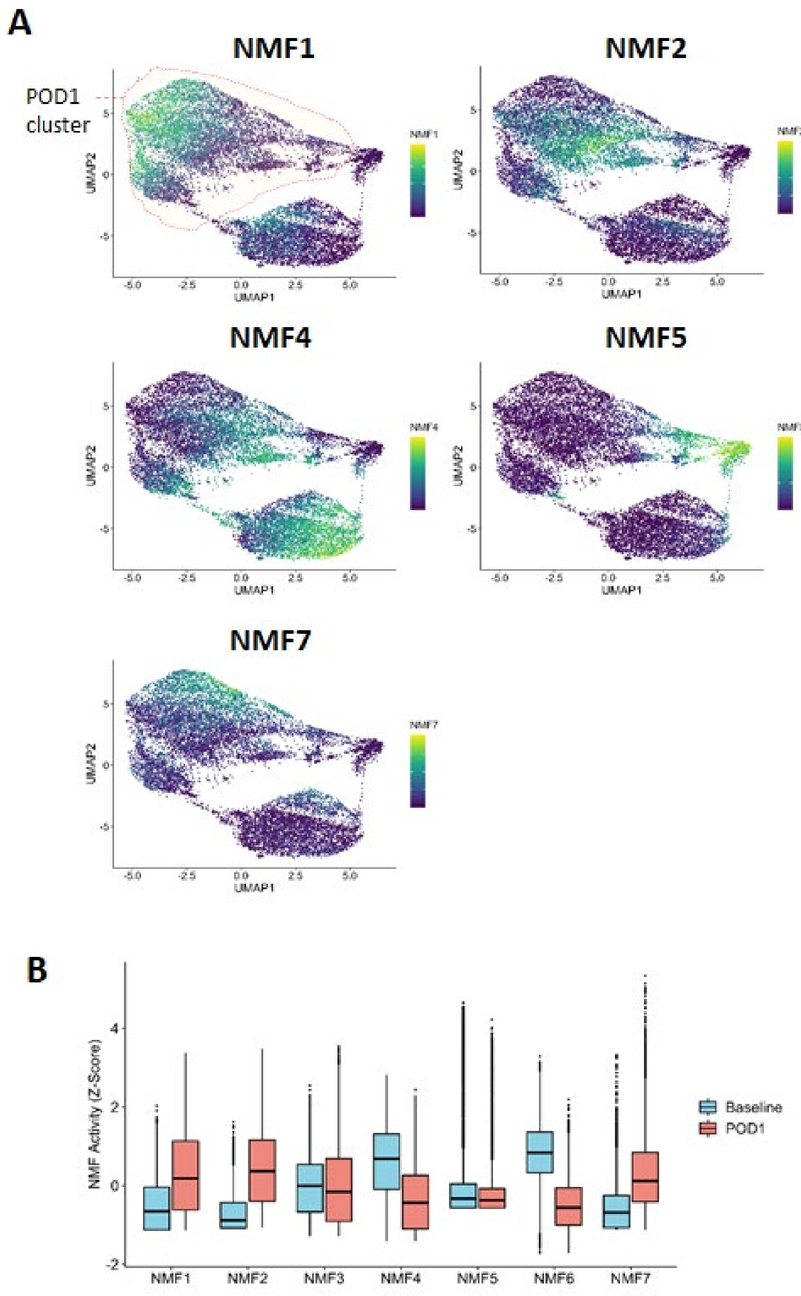


**Supplemental Figure S5. scRNA-seq extended data 2 – NMF plots.**

**(A)** NMF plots of all 7 gene expression profiles and which monocytic cells express them. The highlighted, red circled population in NMF1 demarcates where the POD1 monocytes clustered to help with the interpretation. **(B)** Histograms displaying the number cells that express a certain NMF program either on Baseline or POD1.

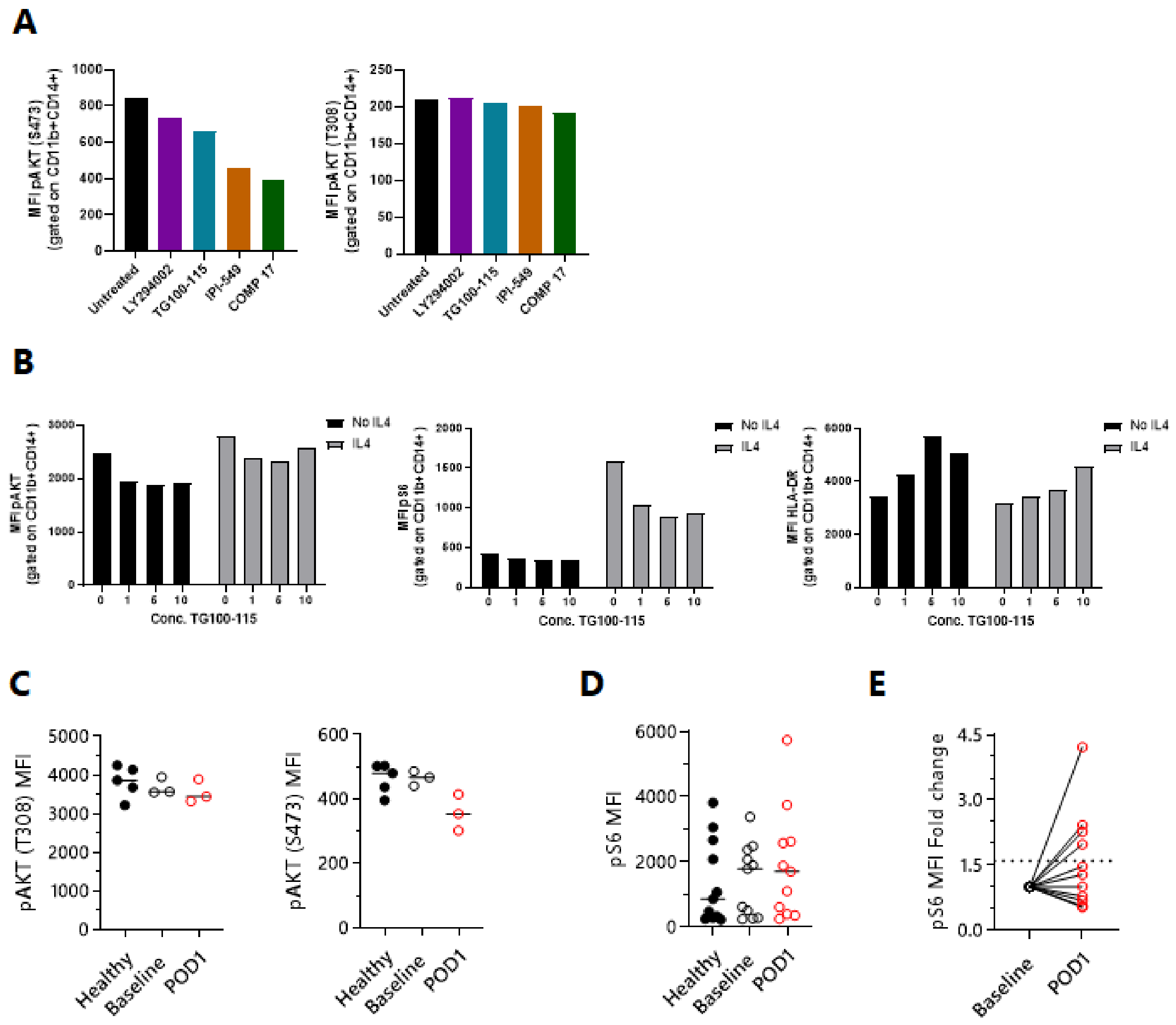


**Supplemental Figure S6. Phospho-flow cytometry for pAKT and pS6 gated on sx-MDSCs +/- PI3K inhibitors.**

**(A)** Effect of PI3K inhibitors (each at 10μM for 40 minutes) on AKT phosphorylation (S473 and T308) in unstimulated whole blood gated on sx-MDSCs (n=1). **(B)** Dose response of TG100-115 on HLA-DR (left), pAKT (mid) and pS6 (right) gated on sx-MDSCs (n=1). **(C)** AKT phosphorylation at T308 (right) and S473 (left) were measured from healthy donors (n=5), and matched cancer surgery patients at Baseline and POD1 (n=3). **(D)** pS6 MFI and **(E)** Fold change in pS6.

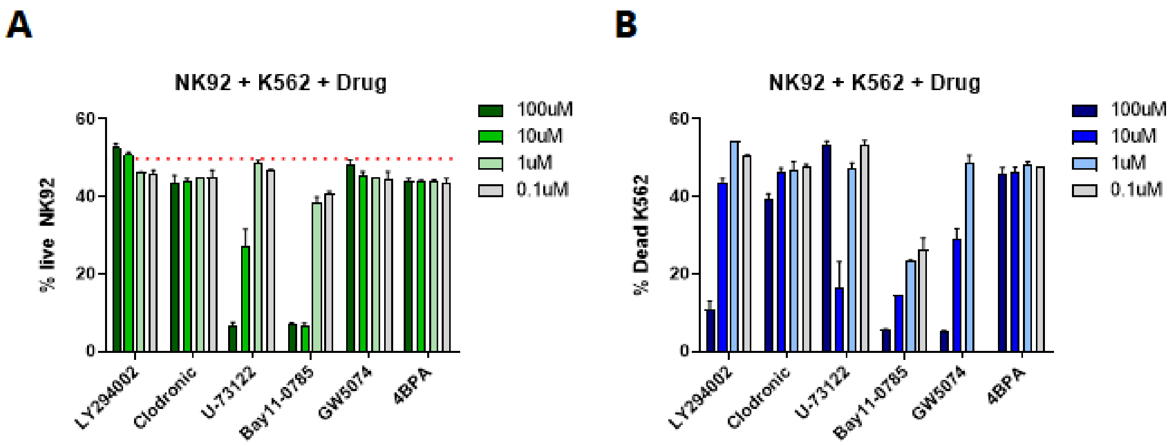


**Supplemental Figure S7. Validation of top candidate compounds following compound screen.**

NK92s were incubated for 24 hours with increasing concentrations of inhibitors. **(A)** Effect of inhibitors on NK cell viability and **(B)** NK cell cytotoxicity.

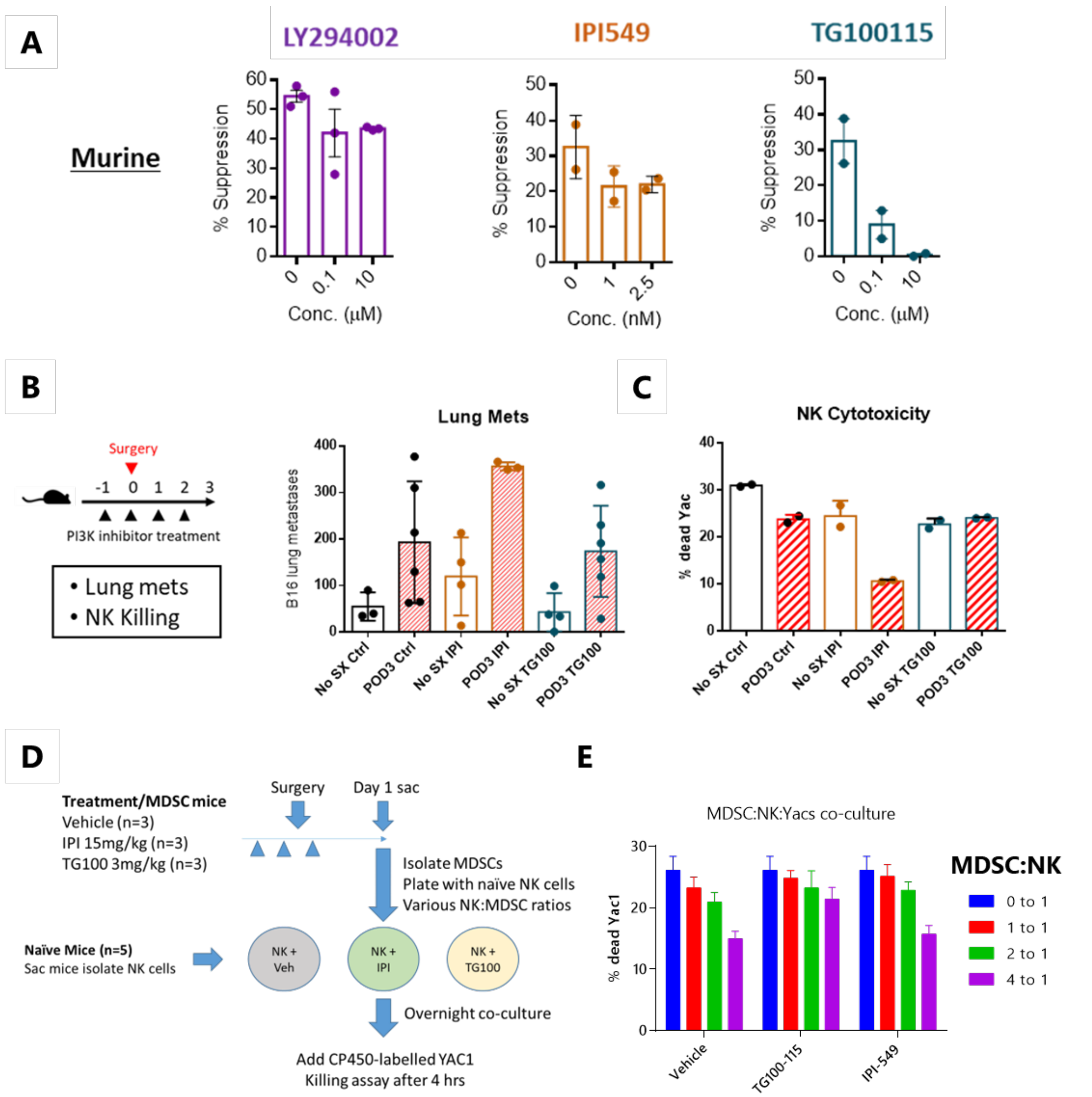


**Supplemental Figure S8. Systemic delivery of PI3K-γ inhibitors does not prevent metastases in our B16F10LacZ murine model of surgical stress.**
**(A)** Ex vivo MDSC:NK suppression assay with murine POD1 MDSCs and naïve NK cells +/- PI3K inhibitors. **(B)** Outline and experimental endpoints (right). Lung metastases after surgery +/- inhibitors (right). **(C)** NK cell function on POD3 +/- inhibitors. **(D)** Schematic of murine MDSC suppression assay. MDSCs were isolated on POD1 from mice following in vivo treatment with PI3K inhibitors and seeded with naïve murine NK cells. **(E)** Results of suppression assay.
